# Perceptual, metacognitive, and computational signatures of temporal landmark uncertainty in tactile motion perception

**DOI:** 10.64898/2026.06.29.735212

**Authors:** Mehdi Adibi

## Abstract

Motion perception depends on estimating the relative timing of sensory events under internal uncertainty. Although perceptual uncertainty is commonly represented by a single internal noise parameter, its structure, sources, and temporal organisation remain poorly understood. Here, I investigated the computational structure of internal uncertainty in phase-based tactile motion perception using continuous amplitude-modulated vibrations delivered simultaneously to two fingertips. Motion discrimination accuracy, response latency, confidence, and confidence entropy exhibited systematic phase-dependent changes, revealing distinct behavioural signatures of temporal uncertainty. Computational analyses demonstrated that tactile motion perception is not explained by a single source of landmark timing uncertainty. Instead, behavioural performance was best accounted for by two additive uncertainty components: an amplitude-dependent component associated with extracting temporal landmarks from the vibration envelope, and an amplitude-independent component shared across stimulus conditions. This dual uncertainty framework consistently explained the frequency dependence of motion perception, the reduced uncertainty observed for sharper vibration envelopes, and the previously reported enhancement of motion perception with exponential compared with sinusoidal modulation. An independent temporal order judgement experiment further validated the uncertainty parameter inferred from motion discrimination, demonstrating that sharpening temporal landmarks reduced timing uncertainty by 35%. Finally, manipulating the initial stimulus state showed that perceptual choices followed the temporal sequence and correspondence of landmark events rather than the initial evolution of the vibration envelopes, providing independent support for landmark-based computations. these findings demonstrate that tactile motion perception is governed by multiple computational sources of landmark timing uncertainty and establish phase-based tactile motion as a tractable paradigm for independently measuring, manipulating, and modelling the computational structure of perceptual uncertainty underlying sensory decisions.

## Introduction

Perception is fundamentally constrained by internal uncertainty. Sensory systems infer external events from noisy, incomplete, and temporally variable signals, and perceptual decisions depend not only on the strength of sensory evidence but also on the structure of the uncertainty associated with that evidence [1 – 4]. Although psychophysical models often represent uncertainty with a single noise parameter, sensory uncertainty can arise from multiple sources, including peripheral encoding, temporal comparison, correlated neural variability, memory of recent sensory samples, and decision processes [5–9]. Disentangling these components is difficult because the latent uncertainty that limits perception is rarely accessible for direct experimental manipulation or independent measurement.

Tactile motion perception provides a powerful setting for addressing this problem. During object manipulation and surface exploration, motion-related tactile cues provide information about contact events, object geometry, texture, friction, and material structure [10–14]. Tactile motion has been studied using moving surfaces, skin stretch, apparent motion, and spatiotemporal sequences of pulsed stimulation across the skin [14–26]. More recent work has shown that temporally structured vibrotactile signals delivered to static skin locations can also support precise temporal, spatial, and cross-digit judgements, including vibrotactile timing judgements, inter-finger phase-shift detection, tactile random-dot motion discrimination, and feature-dependent vibration perception [27–33]. These findings indicate that tactile motion perception is not determined solely by physical displacement across the skin, but can also arise from temporal relationships between spatially separated tactile signals.

In previous work, I developed the physical and computational foundation for phase-based tactile motion perception, showing that robust motion percepts can be generated by delivering two continuous amplitude-modulated vibrations to separate fingertips with controlled phase differences between their envelopes [34]. This framework proposed that motion direction can be inferred from the relative timing of salient temporal features in the vibrotactile envelopes, such as peaks, troughs, or points of maximal envelope change. In the landmark-based account, perceptual errors arise from uncertainty in both the perceived timing and the correspondence of these landmarks across stimulation cycles. The model explains poor accuracy when corresponding landmarks are nearly simultaneous, and also explains ambiguity near anti-phase stimulation, where landmark assignment to the same or neighbouring cycles becomes uncertain. Subsequent psychophysical experiments provided behavioural support for this framework, showing that motion discrimination is largely insensitive to modulation amplitude while envelope pattern and spatial configuration systematically modulate performance and response time [35]. These studies established phase-based vibrotactile stimulation as a controlled paradigm for probing tactile motion inference and perceptual decision-making. However, the statistical structure, stimulus dependence, and underlying computational components of the landmark timing uncertainty that limits perceptual performance remain unresolved.

The present study uses this phase-based tactile motion paradigm to examine the structure of landmark timing uncertainty. The central question is whether perceptual uncertainty in tactile motion can be explained by a single timing-noise parameter, or whether it reflects multiple uncertainty components. This question is important because internal noise is often treated as independent and stationary, whereas biological sensory systems can exhibit temporally correlated variability, stimulus-dependent precision, and decision-related uncertainty [5–9]. In tactile motion paradigm, these components can be separated more directly because the relevant sensory variable is a controllable temporal relationship between envelope landmarks. The present study combines behavioural, computational, and metacognitive analyses to characterise these uncertainty components. First, I quantify how accuracy, response time, confidence, and confidence entropy vary with phase difference, testing whether ambiguous phase relationships produce convergent signatures of perceptual uncertainty [36–40]. Second, I extend the landmark framework from uniform to Gaussian timing uncertainty and test whether correlated landmark noise can account for the observed behavioural structure. Third, I examine whether frequency-dependent changes in performance are better explained by a single timing-noise parameter or by dual uncertainty components. Finally, I directly measure landmark timing precision in an independent temporal order judgement task, showing that sharper envelope peaks reduce the estimated timing uncertainty. Together, these analyses establish phase-based tactile motion perception as a model system for studying how stimulus geometry, temporal landmark extraction, correlated internal noise, confidence, and perceptual decisions are linked through a common uncertainty framework.

## Materials and methods

Psychophysical experiments were conducted to investigate the computational structure of temporal uncertainty underlying phase-based tactile motion perception. The study comprised two complementary behavioural paradigms. The first used a two-alternative forced-choice (2AFC) motion direction discrimination task, following the general procedure described previously [34, 35]. The second employed an independent temporal order judgement (TOJ) task to obtain direct behavioural estimates of landmark timing uncertainty. The motion-discrimination data were obtained using the phase-based vibrotactile motion task described previously [34] and was reanalysed here to characterise the structure of landmark timing uncertainty through accuracy, response time, confidence, and computational modelling. In both tasks, participants completed self-paced discrete trials in which vibrotactile stimuli were delivered to the middle and/or index fingertips of the right hand, and experimental conditions were pseudo-randomised within each session. All experimental procedures were approved by the Monash University Human Research Ethics Committee (MUHREC) (Project ID 27649; date of approval: 2 August 2021) and conducted in accordance with the Declaration of Helsinki.

### Participants

A total of 35 healthy adults (18 female; age range 16–49 years; one left-handed) participated in the study. Twenty-five participants (12 female; age range 19–34 years; one left-handed) completed the motion direction discrimination task [34], and an independent cohort of 10 participants (6 female; mean age 24.1 years, s.d. 9.4 years; all right-handed) completed the temporal order judgement (TOJ) experiment. Participants were recruited from Monash University (*n* = 33 undergraduate and graduate students) and the surrounding community. All participants reported normal tactile perception and no history of neurological disorders, and provided written informed consent prior to participation.

### Apparatus and Vibrotactile Stimulation

The apparatus and stimulation procedures were identical to those described previously [34, 35]. Briefly, vibrotactile stimuli were delivered simultaneously to fingertips using miniature solenoid transducers (PMT-20N12AL04-04, Tymphany HK Ltd; 4 Ω, 1 W, 20 mm diameter) mounted 5 cm apart on a vibration-isolated platform. Stimuli were generated in MATLAB (MathWorks Inc.) at a sampling rate of 44.1, 48 or 192 kHz, and delivered through either a Creative Sound Blaster Audigy Fx 5.1 sound card (model SB1570) or a Creative Sound Blaster Live! 24-bit External (model SB0490). The shape and curvature of the transducer matched the size and contour of adult fingertips [41]. In the TOJ experiment, actuators were driven using a 3 W stereo amplifier module (PAM8403 chipset). Across experiments, actuator outputs were matched across channels and configurations. Sound pressure levels (SPL) were measured 5 mm from each actuator and adjusted to produce matched outputs (mean SPL = 78.9 dB). Matching was further verified perceptually to ensure comparable stimulus intensity across actuators.

Vibrotactile stimuli consisted of a sinusoidal carrier (*f*_*c*_ = 100*Hz*), with the amplitude modulation, spatial configuration, and temporal parameters depended on the specific experimental task, as described below. Although the carrier frequency falls within the audible range, pilot testing confirmed that the stimuli were not audible to participants and were perceived only through touch [34]. Across all experiments, the modulation amplitude was set well above detection threshold. Additionally, Previous work further demonstrated that, once above detection threshold, motion discrimination was largely invariant to modulation amplitude, with even a twofold reduction producing negligible effects on behavioural performance [35].

### Motion direction discrimination task

Participants performed a discrete-trial two-alternative forced-choice (2AFC) task to judge the perceived direction of vibrotactile motion. On each trial, two amplitude-modulated vibrations were delivered simultaneously to the index and middle fingertips of the right hand. Participants gently rested their fingertips on the transducers without applying force while maintaining a stable hand posture throughout the experiment (Figure 1A). The transducers were mounted 5 cm apart on a vibration-isolated platform such that vibrations from one actuator were not perceptible at the other. Participants reported clear perception of the envelope modulation.

**Figure 1.**
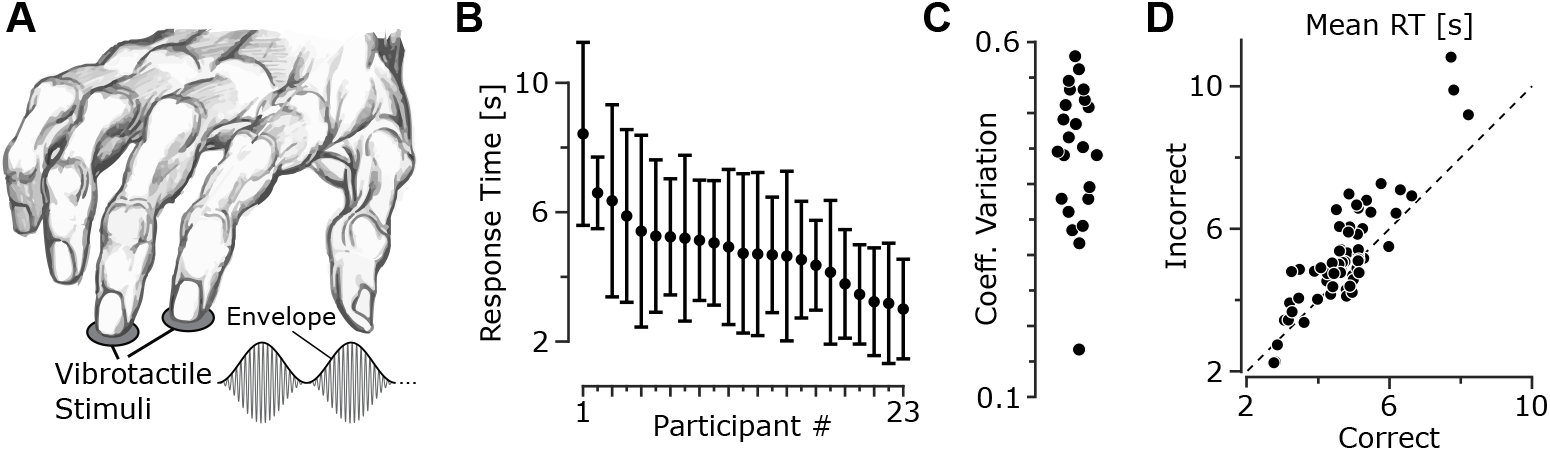
Inter-subject variability in the motion direction discrimination task. **A**, Motion discrimination task. Participants received two AM vibrotactile stimuli with varying phase differences in their index and middle fingerpads on every trial. **B**, Net mean and standard deviation (s.d.) in response times (RTs) per subject across all trials. Subjects are sorted based on their mean RT in descending order. Error bars indicate s.d.. **C**, The RT coefficient of variation (CV). Markers represent individual subjects. **D**, The mean RT for incorrect trials vs. correct trials. Markers represent individual subject and stimulus pairs.

Each trial consisted of three modulation cycles (e.g., 6 s at an envelope frequency of 0.5 Hz). For a phase difference Δ*φ*, the two sinusoidally amplitude-modulated vibrations were

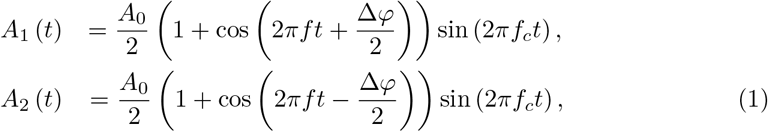

where *f* and *f*_*c*_ denote the envelope and carrier frequencies, respectively. The peak-to-peak amplitude of the output waveform was set to 1.98 V.

To eliminate directional cues arising from the from differences in initial envelope amplitude, stimulus onset was chosen randomly from one of the two iso-amplitude points of the modulation cycle. For non-zero Δ*φ*, these occur at *t*_0_ = 0 and 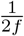, corresponding to envelope amplitudes of 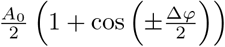 and 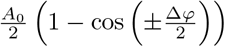, respectively (Figure 6A). For a zero phase offset, the same onset conditions (*t*_0_ = 0 and 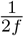) were used. These two points correspond to the maximum possible difference between the two onset amplitudes across all tested phase offsets (Figure 6B), despite the two vibrations remaining perfectly in phase throughout the trial. At 180°, the two onset conditions have identical envelope amplitudes throughout stimulus onset and differ only in the direction of the initial envelope change preceding the first landmark peak. For consistency across phase differences, these onset conditions are referred to as the high and low initial-state conditions. The two onset conditions were presented with equal probability on each trial.

Participants indicated the perceived direction of motion (leftward or rightward) by pressing the corresponding arrow key with their left hand. Responses were self-paced and could be made at any time during or after stimulus presentation.

The motion discrimination dataset was based on the experimental paradigm described previously [34] with a total of 11,903 trials. Sinusoidal envelopes were presented with phase differences ranging from -180° to 180° (30° increments) at modulation frequencies of 0.5, 1 and 1.5 Hz. A subset of participants completed the task at a fixed modulation frequency of 0.5 Hz using phase differences of 0°, ±30°, ±60°, ±90° and 180°, in which confidence ratings were collected after each trial using a five-point scale. Confidence ratings were linearly transformed to a 0–100% scale for analysis. Experimental conditions were presented in pseudorandom order, with approximately 30 repetitions per condition.

### Tactile Landmark Temporal Order Judgement Task

To quantify the temporal precision of tactile landmark detection, ten participants (8 female; all right-handed; age 24.1 ± 9.4 s.d. years, range 16–49 years) performed a two-alternative forced-choice (2AFC) temporal order judgement (TOJ) task. On each trial, participants judged whether a brief auditory probe occurred before or after the perceived peak of a vibrotactile stimulus delivered to the middle fingertip of the right hand.

Vibrotactile stimulation was delivered to the middle fingertip of the right hand. Each stimulus consisted of a single modulation cycle (2 s duration; 0.5 Hz envelope frequency; 100 Hz carrier). Two envelope shapes were tested: a broad sinusoidal envelope (cos^1^) and a sharper envelope generated by raising the sinusoid to the fifth power (cos^5^). Both envelopes reached the same peak amplitude and contained a single well-defined envelope peak.

The auditory probe was a 10 ms tone at 1 KHz presented binaurally through headphones. On each trial, the probe occurred at one of three temporal separations 50, 150, or 300 ms relative to the tactile envelope peak. For each separation, the probe was presented either before or after the tactile peak with equal probability, yielding signed temporal offsets of ±50, ±150, and ±300 ms. Participants indicated whether the auditory probe occurred before or after the perceived tactile peak using two foot switches. Each participant completed 50 trials for every combination of envelope shape, temporal separation, and temporal order, resulting in 300 trials per participant and a total of 3,000 trials across the experiment.

### Psychometric Modelling

To estimate landmark timing uncertainty, trial-level responses were analysed using generalised linear fixed-effects models with a binomial distribution and probit link function. The models were fit to the psychometric data, defined as the likelihood of reporting the auditory probe as occurring after the tactile landmark, as a function of the signed temporal offset between the auditory probe and tactile landmark. Under the Gaussian uncertainty assumption of the landmark framework, the probit slope provides a direct estimate of the landmark timing uncertainty parameter, *σ*. Several candidate models were considered, including fixed-effects models with and without an intercept term, as well as models incorporating subject-specific random intercepts and random slopes. Model selection was based on Akaike Information Criterion (AIC), Bayesian Information Criterion (BIC), and maximised log-likelihood. The best-fit model included temporal offset and the interaction between temporal offset and envelope shape as fixed effects, without subject-level random effects. This model substantially outperformed all alternative formulations (ΔAIC ≥ 232, ΔBIC ≥ 238). The absence of an intercept constrains choice likelihood to chance level (50%) at zero temporal offset, consistent with the symmetry of the task and the assumptions of the landmark framework. The estimated dispersion parameter of 1.00 indicated no evidence of deviation from the binomial assumption.

Under the Gaussian uncertainty assumption of the landmark framework (probit link), the timing uncertainty parameter *σ* was estimated as *σ* = 1*/β*, where *β* denotes the fitted probit slope parameter. The mean of *σ* was estimated using a second-order Taylor expansion about the fitted value of *β*, and standard errors (s.e.) were obtained using first-order error propagation (delta method). Given the fitted values o *β* and their relatively small standard errors, the second-order correction term was negligible, and the reported estimates were effectively identical to *σ* = 1*/β*.

To quantify the effects of envelope shape on temporal-order discrimination accuracy, trial-level responses were analysed using a generalised linear mixed-effects model with a binomial distribution and logit link function. Accuracy was modelled as a function of the absolute temporal offset between the auditory probe and tactile landmark and its interaction with envelope shape. Subject-specific random slopes were included for temporal offset to account for individual differences in temporal sensitivity. The model was fit without an intercept term, and the effect of envelope shape was included only through its interaction with temporal offset in both the fixed and random effects. This parameterisation constrains accuracy to chance level (50%) at zero temporal offset. The constraint was based on the symmetry of the task, and was consistent with the preceding probit analysis of choice likelihood, in which model selection favoured a no-intercept psychometric function symmetric around zero temporal offset. The logit model is based on the well-established logistic psychometric function [42]. Model diagnostics indicated no evidence of overdispersion relative to the binomial assumption (dispersion parameter = 0.990).

### Probabilistic model of landmark perception

For an AM vibration with envelope *A* (*t*), sensory noise and perceptual limits were assumed to introduce uncertainty in the perceived timing of the envelope peak. The perceived peak time was modelled as a normally distributed random variable centred on the true peak, with standard deviation *σ*_amp_. This amplitude-dependent uncertainty was derived from the temporal window around the peak over which the envelope remained within an amplitude threshold *δ*, defined by |1 − *A* (*t*) */A*_0_| ≤ *δ*, where *A*_0_ denotes the peak envelope amplitude.

For the envelope used in the TOJ task,

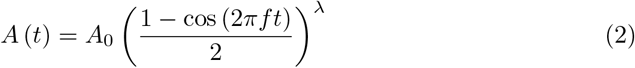

where *f* = 0.5 Hz and *λ* = 1 or 5, the width of the near-peak uncertainty window is:

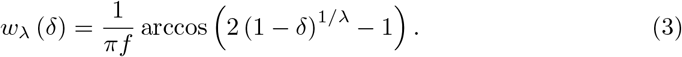

This window can be converted to an equivalent Gaussian uncertainty in several ways. More generally, if the interval is assumed to contain a fraction *q* of the Gaussian probability mass, the corresponding standard deviation (s.d.) is

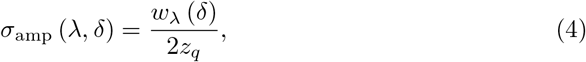

where *z*_*q*_ is the Gaussian quantile corresponding to the central probability mass *q* (e.g., *z*_*q*_ = 1.96 for a window containing 95% of the landmark uncertainty, i.e., *q* = 0.95). Here, for consistency with the uniform formulation used in our previous work [34], the uncertainty window was converted to an equivalent Gaussian uncertainty by matching the variance of a uniform distribution over the same interval, corresponding to 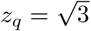, yielding

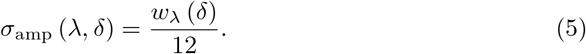

To account for uncertainty sources independent of envelope shape, a shared additive uncertainty component *σ*_0_ was introduced in variance form:

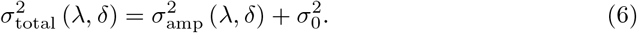

The threshold parameter *δ* and shared uncertainty component *σ*_0_ were estimated by solving the system of equations using uncertainty estimates from the hierarchical probit psychometric analysis, for the broad and sharp envelope conditions:

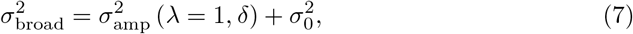

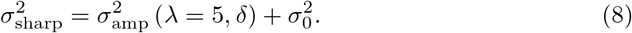

Model performance was assessed by comparing the resulting psychometric predictions with the observed mean accuracy across temporal offsets.

### Probabilistic landmark model of tactile motion perception

Consider two AM vibrations with phase difference Δ*φ*, which corresponds to a temporal lag *d* between their envelopes:

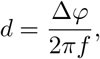

where *f* is the envelope modulation frequency. Let *t*_1_ and *t*_3_ denote the perceived moments of two successive salient landmarks (e.g., peaks) of vibration 1, and let *t*_2_ denote the corresponding landmark of vibration 2 that occurs between *t*_1_ and *t*_3_. These landmarks are extracted from the envelopes of the AM vibrations, and their relative timing determines the motion direction, as follows. Since both vibrations are periodic with identical envelope shapes (with vibration 2 being a phase-shifted version of vibration 1), the landmark of vibration 2 is offset by *d*. Likewise, *t*_3_ is one cycle after *t*_1_, with an offset of envelope modulation period 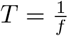. Without loss of generality, we assume *d >* 0, and align the salient landmarks relative to zero.

Hereafter, we focus on peak features, though the same logic generalises to other salient envelope landmarks such as troughs or moments of maximum envelope derivatives (changes). Due to sensory noise and perceptual uncertainty, the detected peaks **t** = [*t*_1_, *t*_2_, *t*_3_]^T^ are assumed to follow a multivariate Gaussian distribution:

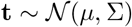

with mean vector *µ* = [0, *d, T* ]^T^ and the covariance matrix Σ. Owing to the periodic and symmetric structure of the stimulus, we assume the marginal variances (i.e., the diagonal of Σ) are identical, denoted by *σ*^2^.

Correct perception of motion direction involves two forms of perceptual computations: first, the **temporal order judgement**, i.e. whether the peak of vibration 2 occurs after that of vibration 1 (i.e., *t*_2_ *> t*_1_). Second, the **inter-peak interval discrimination**, i.e., whether the time interval from *t*_1_ to *t*_2_ is shorter than from *t*_2_ to *t*_3_, i.e., *t*_2_ − *t*_1_ *< t*_3_ − *t*_2_. To analyse both jointly, define the auxiliary transformed random vector **u** = [*u*_1_, *u*_2_]^T^ derived from the **t** via the linear transformation:

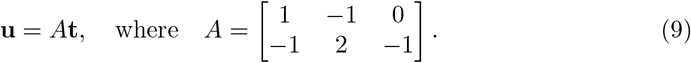

Here, *u*_1_ = *t*_1_ − *t*_2_ and *u*_2_ = 2*t*_2_ − *t*_1_ − *t*_3_ represent the two decision variables. Since **t** is Gaussian, so is **u**:

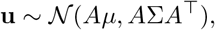

with mean:

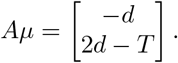

### Structure of covariation

The perceived time of one landmark may covary with the perceived time of others, particularly if they are temporally proximal. This interdependence may arise from shared sensory encoding processes or central inference mechanisms. To capture this, we model the covariance matrix Σ with an exponentially decaying correlation structure, analogous to the Ornstein–Uhlenbeck (OU) process. Specifically, the correlation between any two landmarks is modelled as:

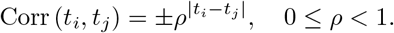

The sign may be positive or negative, depending on whether the peaks bias each other toward (positive) or away from (negative) each other. This differs from the standard OU process, which only produces positive correlations. Here, I adopt this flexible structure to allow for either scenario. Under this model, the covariance matrix Σ takes the form:

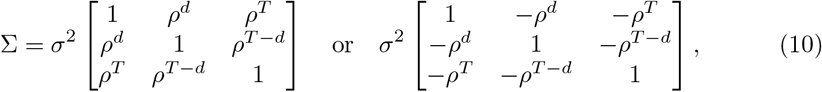

This formulation ensures temporal stationarity, meaning that the correlation between perceived landmark times depends only on their temporal separation. The special case of *ρ* = 0 corresponds to uncorrelated (and hence independent due to normality) uncertainties across landmarks.

The covariance matrix of the transformed variables **u** is then:

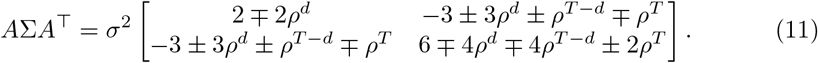

This bivariate normal distribution over **u** characterises the joint uncertainty in temporal order and interval discrimination, allowing us to compute the probability of a correct motion judgement analytically.

### Temporal order judgement

The first source of error arises from uncertainty in judging the relative timing of salient landmarks (e.g., peaks) across the two vibrations. Specifically, the probability that the landmark from envelope 1 is perceived before that of envelope 2 corresponds to the cumulative distribution function (c.d.f.) of the difference in perceived times, *u*_1_ = *t*_1_ − *t*_2_, evaluated at zero.

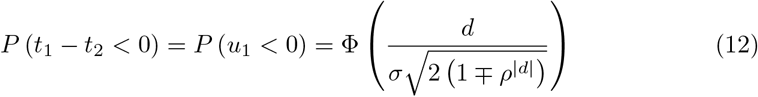

where Φ( ) denotes the standard c.d.f.. The ∓ sign reflects the effect of positive versus negative correlation on the variance of the difference..

### Cross-cycle ambiguity in inter-peak interval discrimination

As the time lag *d* increases, the uncertainty in estimating *t*_2_ makes it increasingly likely that its perceived timing falls closer to the *next* peak of envelope 1 at *t*_3_ rather than the original one at *t*_1_. This cross-cycle ambiguity results in an incorrect interval comparison: the perceived delay from *t*_1_ to *t*_2_ exceeds that from *t*_2_ to *t*_3_, i.e., *t*_2_ − *t*_1_ *> t*_3_ − *t*_2_, or equivalently, *u*_2_ *>* 0, thus misjudging the motion direction. The probability of correctly assigning *t*_2_ to the current modulation cycle is given by the c.d.f. of *u*_2_, evaluated at zero:

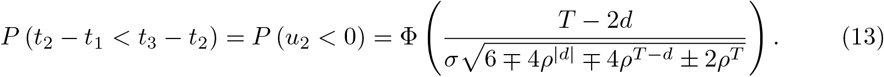

### Joint probability of correct motion perception

Correct motion perception requires both correct temporal order identification of peaks and correct cross-cycle inter-peak interval discrimination, i.e., **u** *<* 0. These two conditions are not independent, and their joint probability must be evaluated accordingly. Here, I compute this probability conditionally, as follows:

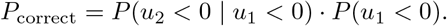

Using the law of total probability over the marginal distribution of *u*_1_, the conditional term can be rewritten as:

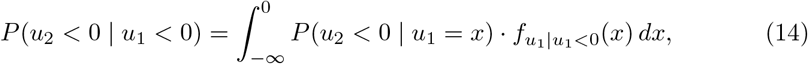

where 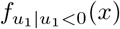 is the conditional probability density function (p.d.f.) of *u*_1_ given *u*_1_ *<* 0. Since Since 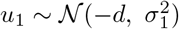, this conditional p.d.f. is given by:

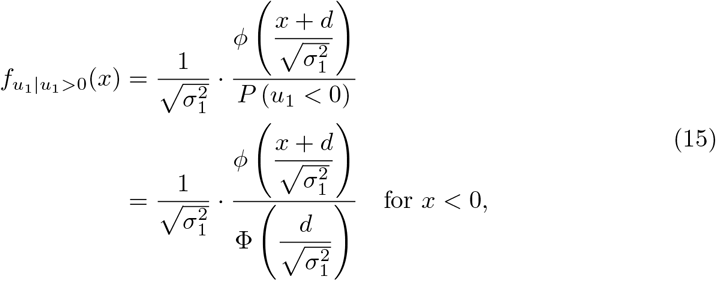

where *ϕ*( ) is the standard normal p.d.f., 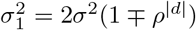, and the normalisation constant *P* (*u*_1_ *<* 0), as given in Eq. 12.

To compute *P* (*u*_2_ *<* 0 | *u*_1_ = *x*), I use a direct substitution method that respects the generative structure. Conditioning on *u*_1_ = *x* implies *t*_1_ = *x* + *t*_2_, which substituted into *u*_2_ = 2*t*_2_ − *t*_1_ − *t*_3_ gives:

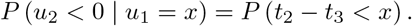

Given 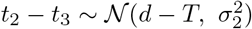 with 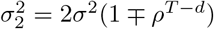, the conditional probability becomes:

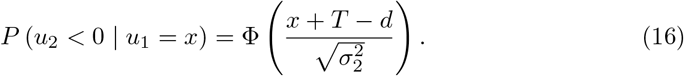

Note that knowing that *u*_1_ = *x* alters the variance, though the new variance does not depend on the specific value of *x*, and shifts the mean by (*x − d*). Putting all terms together, the joint probability of correct perception is:

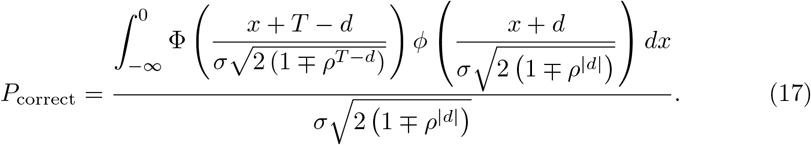

For the special case when *t*_1_, *t*_2_ and *t*_3_ are independent, i.e., when *ρ* = 0, the expression above becomes:

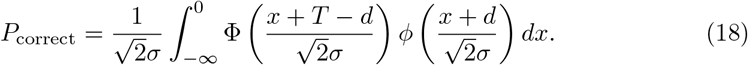

### Robustness of the tactile motion model to noise distribution

To quantify the influence of the sensory noise distribution on the tactile motion model predictions, I computed the extent to which the shape of noise (i.e., Gaussian versus uniform) affects the predicted probability of correct motion perception. For both distributions, the standard deviation *σ* was matched. The divergence in predictions was quantified as the mean absolute difference between the two models’ predictions, normalised by their mean performance, defined as:

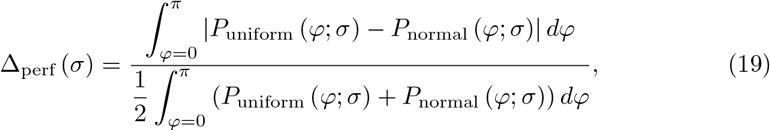

where *P*_normal_ (*φ*; *σ*) denotes the predicted probability of correct motion direction perception at phase difference *φ* under Gaussian timing noise, as defined in Eq. 18, and *P*_uniform_ (*φ*; *σ*) denotes the corresponding prediction under a uniform noise model, derived analytically in closed form in [34]. The integrals were evaluated numerically across phase differences *φ* in 1° increments.

To relate these differences in predicted behaviour to the underlying difference in noise distributions, I also computed the total variation distance between the two probability density functions (p.d.f.s), defined as:

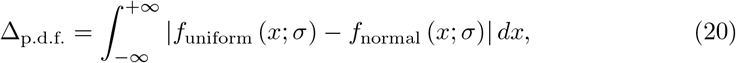

where *f* (*x*; *σ*) denotes the p.d.f. of a distribution with identical standard deviation *σ* and mean. This expression after change of variables is independent of the mean, and independent of *σ* as well (see below). Note that Δ_p.d.f._ has the same form as Δ_perf_ normalised to the mean of the two distributions, but since both p.d.f.s integrate to 1, the denominator simplifies to 1.

Assuming a mean of zero (without loss of generality), the p.d.f. of a uniform distribution with standard deviation *σ* is constant with the value of 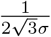 within the range 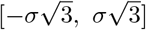, and zero outside. Therefore, the total variation distance becomes:

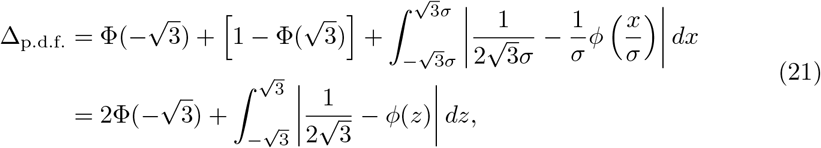

where *z* = *x/σ* is the normalised variable. This integral can be expressed in terms of the crossover point *z*^∗^ where 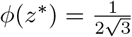, which solves:

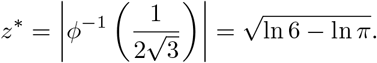

where *ϕ*^—1^ (·) denotes the inverse of the standard normal distribution. Using symmetry and integration over the appropriate regions:

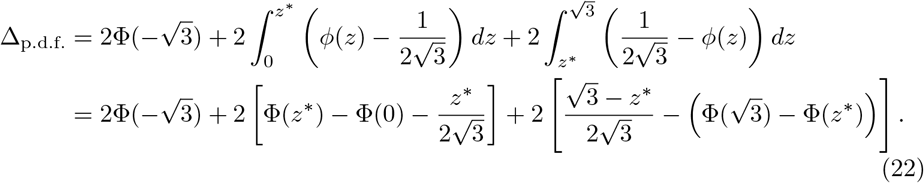

Evaluating the expressions numerically yields:

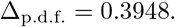

This value reflects the absolute divergence in shape between the two noise distributions, and can be used to interpret the behavioural-level divergence Δ_perf_(*σ*), allowing comparison between low-level noise structure and perceptual predictions.

### Cross-Validation for Phase-Level Generalisation

To assess model generalisability to unseen phase differences, I used a leave-one-out approach. For each of the tested phase differences (30° to 150°), the model was fit to the remaining phase values across all frequencies. The fitted model was then used to predict performance at the left-out phase across frequencies. This process was repeated for each phase difference. Predictions were computed separately for each subject and averaged. I also compared these predictions to the model predictions when trained on the full dataset including the held-out phase, and the corresponding empirical performance. The r.m.s.e. values and prediction-empirical correlations were computed to quantify accuracy.

## Results

Previous work established phase-based tactile motion perception as a robust paradigm in which directional motion is inferred from temporal phase differences between continuous amplitude-modulated vibrations delivered to two fingertips [34, 35]. Those studies developed the landmark framework for tactile motion perception and demonstrated that motion discrimination depends primarily on temporal phase relationships rather than vibration amplitude, while envelope dynamics and spatial configuration modulate perceptual sensitivity. Here, I use this paradigm (Figure 1A) to investigate the computational structure of landmark timing uncertainty. Specifically, I characterise behavioural signatures of uncertainty through analyses of response accuracy, response time, confidence, and confidence entropy, evaluate alternative models of landmark uncertainty, and independently quantify landmark timing precision using a temporal order judgement task.

### Diverse response behaviour

Participants exhibited substantial inter-individual variability in response times (RTs), with mean RTs ranging from 3.02 s to 8.41 s (Figure 1B). The coefficient of variation (CV) of RTs, a within-subject measure of response variability, ranged from 0.17 to 0.58 (median: 0.45), reflecting marked differences in decision-making speed and consistency across individuals (Figure 1C).

Despite this variability, a consistent pattern was evident when comparing accuracy conditions: across subject–stimulus (phase difference) pairs, incorrect trials were slower than correct ones by an average of 512 ms ± 100 ms (s.e.m.; Figure 1D). A paired t-test across subject-stimulus pair means confirmed that this difference was significant (*p <* 10^—5^). Additionally, 72.1% of these pairs showed longer RTs for incorrect responses (*p <* 0.001, Fisher’s exact binomial test; ). These findings are consistent with prior results [34], suggesting that longer RTs tend to occur under conditions of greater sensory uncertainty.

### Confidence ratings are inversely related to RT and modulated by accuracy

Confidence ratings indexed participants’ subjective perceptual certainty providing a metacognitive index of how strongly they perceived the motion signals on each trial [34]. Similarly, RTs provides a behavioural index of the strength of sensory evidence, and mirror the average confidence rating pattern as a function of phase difference, indicating reduced confidence for ambiguous stimuli [34]. To quantify the relationship between confidence judgements and RTs, I computed Pearson’s correlation between confidence ratings and RTs for each participant. All participants exhibited negative correlations with all but one being statistically significant (mean *r* = −0.33 ± 0.05 s.e.; all *p <* 10^—4^, one non-significant at *p* = 0.29). Correlations were averaged in Fisher *z* space and transformed back for reporting, and s.e. was estimated using the delta method (see Methods). The inverse relationship remained when accounting for accuracy via partial correlation (mean *r*_partial_ = 0.24 0.04; all but one *p <* 0.05), confirming that slower responses were associated with lower confidence, regardless of correctness (Figure 1D).

To directly characterise the relationship, I fitted linear mixed-effects models predicting confidence ratings (0–100%) from standardised RT (*z*-scored within subject), accuracy (correct vs. incorrect), and their interaction. A random intercept was included for subject to account for individual differences. Model comparison confirmed that including an interaction term between RT and accuracy significantly improved model fit (ΔAIC = 6.01, *χ*^2^(1) = 7.41, *p* = 0.0065), compared to a model without the interaction. The final model estimates showed that confidence was significantly higher on correct trials (*β* = 7.58, 95% CI [6.01, 9.15], *p <* 10^−20^), and decreased as RT increased (*β* = −7.29, 95% CI [-8.41, -6.16], *p <* 10^−36^), as shown in Figure 2A. The interaction term was also significant (*β* = 1.81, 95% CI [0.51, 3.11], *p* = 0.0065), indicating indicating a steeper RT-confidence slope for incorrect trials.

**Figure 2.**
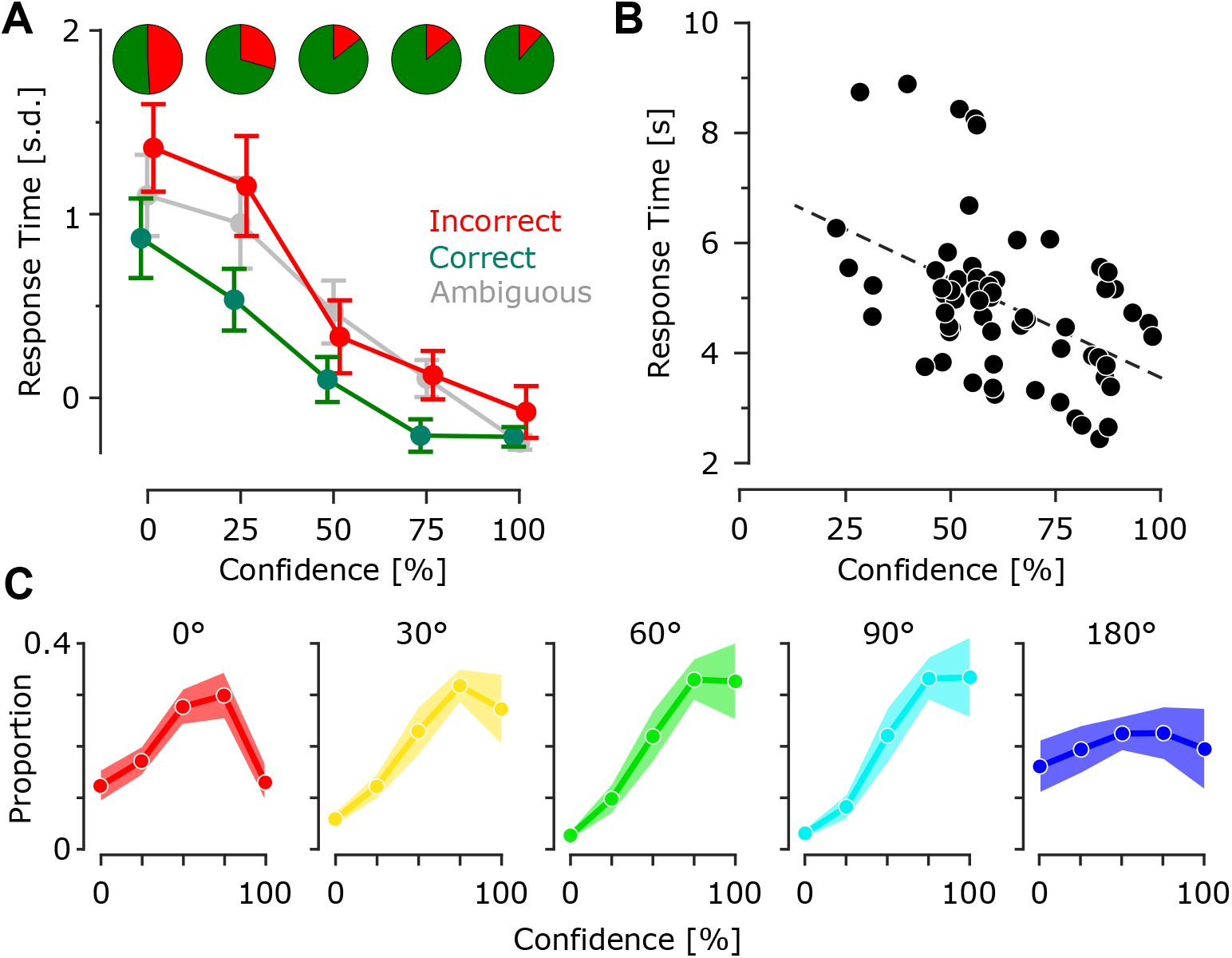
Relationship between confidence, accuracy and RT. **A**, Average standardised RT as a function of confidence ratings for correct (green), incorrect (red) and ambiguous (0° and 180°, grey) trials averaged across subjects. Pie charts indicate proportion of correct and incorrect trials per confidence ratings. **B**, Average RT vs. confidence ratings per subject-stimulus pair. The dashed line represents the linear regression fit (slope: − 0.036, intercept: 7.13 s, *p* = 1.26 × 10^—4^, *R*^2^ = 0.216) **C**, Histogram of confidence ratings per stimulus averaged across subjects. The shades represent s.e.m..

For trials with ambiguous stimuli (0° and 180° phase difference), where accuracy is undefined, a separate linear mixed-effects model predicting confidence solely from RT was fitted, with subject as a random effect. The model revealed a robust negative effect of RT on confidence: confidence decreased by 9.2% per 1 s.d. increase in RT (*β* = − 9.23, 95% CI [-10.26, -8.21], *p <* 10^—64^). Despite the absence of ground-truth correctness in these ambiguous trials, participants’ confidence continued to inversely scale with RTs, mirroring the pattern observed in structured stimuli. Together, these results reveal a consistent and interpretable behavioural pattern: longer decision times are associated with lower subjective confidence, across both defined and ambiguous perceptual conditions. Confidence ratings reflect not only outcome (accuracy) but also internal processing dynamics (RT), suggesting that participants integrate decision latency as a cue for uncertainty. Together, these results reveal a consistent pattern where longer decision times are associated with lower subjective confidence regardless of accuracy, suggesting that confidence and RT reflect a shared underlying decision variable. Despite inter-individual differences in response profiles, the inverse RT-confidence relationship held consistently across subjects.

### More confident subjects are systematically faster

I also observed a significant negative correlation between average reaction time and confidence across subjects, that slower responses were associated with lower confidence (Pearson’s *r* = − 0.46, *p* = 1.26 × 10^—4^; Figure 2B). Unlike the within-subject effects captured by the mixed-effects models above, this across-subject correlation reveals a stable individual trait: participants who were slower overall also tended to be less confident on average. This suggests that inter-individual differences in decision speed may reflect broader differences in perceptual certainty or decision thresholds.

### Phase-dependent variations in perceptual certainty and confidence entropy

To further characterise the perceptual certainty for each phase difference condition, Figure 2C depicts the average distribution of confidence ratings as a function of phase difference. At 0°, confidence ratings were more frequent around mid-range values (50% and 75%), consistent with previous findings that this condition yields perceptually ambiguous motion due to a lack of consistent directional cues [34]. Intermediate phase differences (30°–90°) elicited increasing confidence, with histograms peaking at high ratings (75% and 100%), indicative of stronger sensory evidence. Interestingly, the 180° condition produced a relatively uniform confidence distribution – unlike the 0° condition, which showed a mid-range skew – indicating that this uncertainty may stem not from a lack of sensory evidence (as for 0° with no directional cue), but from internal decision biases or subtle onset asymmetries in the vibrations. Thus, while the mechanisms may differ, both 0° and 180° phase differences elicit similar behavioural markers of perceptual uncertainty: slower RTs, lower confidence, and broader or disperse confidence distributions.

This is further supported by the entropy of the confidence distributions, which was higher for the 0° (1.99±0.06 bits) and 180° (1.79±0.12 bits) phase differences – reflecting a more uniform and less decisive pattern of confidence across trials – and lower at intermediate phase differences: 60° (1.61±0.13 bits) and 90° (1.56 0.15 bits). Entropy provides a quantitative index of perceived uncertainty, capturing not only performance ambiguity (as with RT and accuracy) but also the subjective dispersion in confidence reports themselves. Higher entropy indicates broader, less peaked confidence ratings, consistent with increased metacognitive uncertainty. This metric complements RT and accuracy by indexing second-order uncertainty in the perceptual decision process, and reveals latent ambiguities that might not be visible from accuracy alone.

### Landmark-Based Model of Tactile Motion Perception

Previously, I proposed a computational, mechanistic model of tactile motion perception based on relative timing of local salient features in vobrotactile stimuli across the fingers [34]. In this model, the tactile system detects specific *temporal landmarks* in the envelope of each vibration – such as peaks, troughs, inflection points, moments of high envelope speed or other salient features – and infers motion direction from their relative timing across fingers. To account for perceptual noise in landmark timing, the model posits a threshold-based mechanism, whereby a landmark (e.g., a peak) is detected when the local change in the envelope exceeds a fixed slope or amplitude threshold. Sub-threshold fluctuations are not perceived as distinct events, resulting in a uniform distribution for the perceived timing of these landmarks. This framework captures two principal sources of perceptual error: (1) local temporal ambiguity, when landmarks occur close in time and their uncertainty distributions overlap, and (2) cross-cycle misattribution, when phase differences approach 180°, causing confusion between temporally adjacent cycles. I previously derived closed-form analytical expressions for the model under the assumption of uniform timing uncertainty, enabling predictions of choice probabilities as a function of lag or phase difference [34] (Figure 3A).

**Figure 3.**
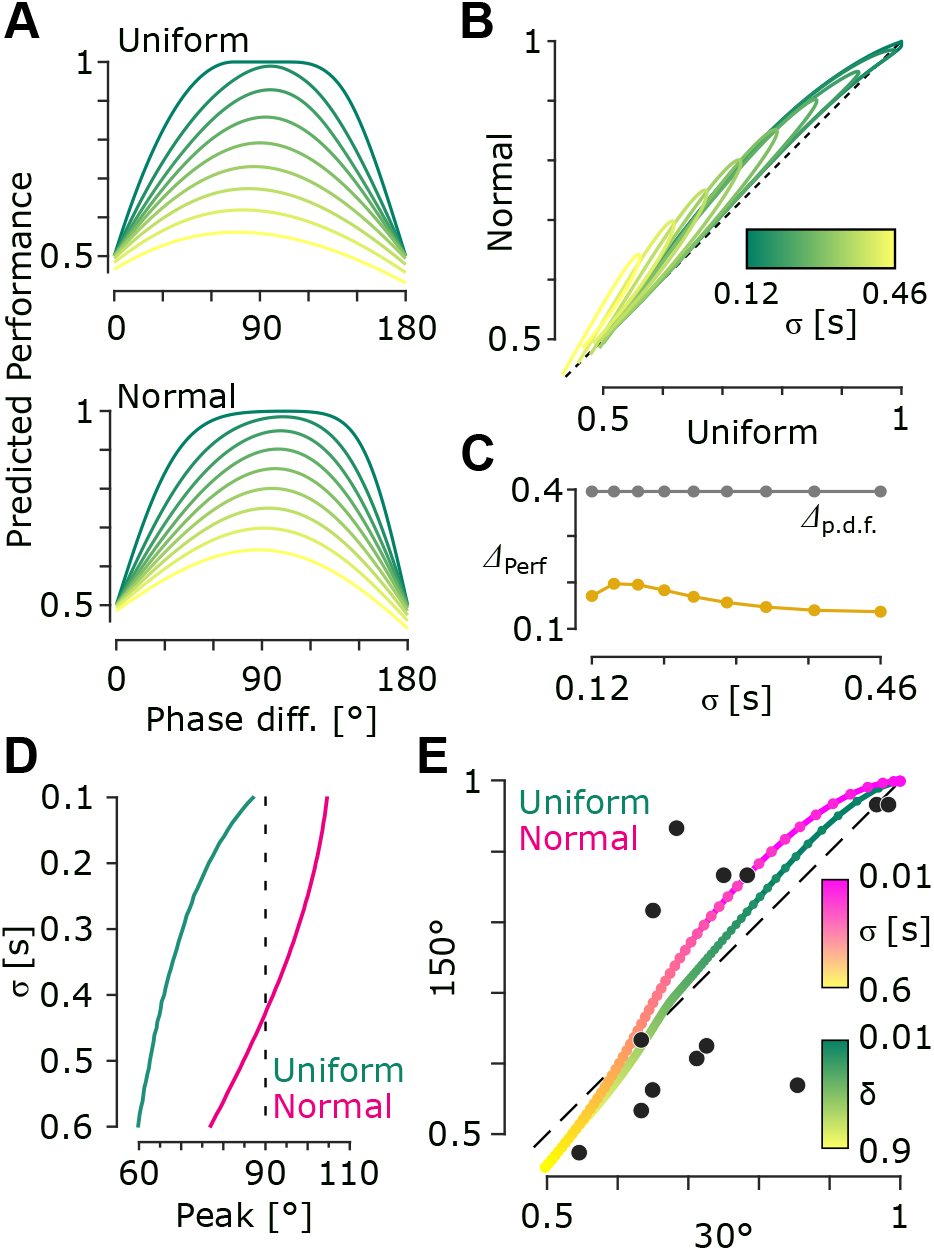
Landmark timing model of motion perception. **A**, The model predictions assuming uniform (top) and Gaussian (bottom) distribution of the internal uncertainty. Curves (shades of green) correspond to 0.1–0.9 envelope amplitude thresholds (relative to envelope peak), in 0.1 increments for uniform, and same variance for Gaussian, equivalent to distribution s.d. of 0.12–0.46 s. **B**, As in **A** but plotting uniform-predicted performances against Gaussian predictions for same phase differences per each variance value. **C**, Net difference in predictions (Δ_perf_, in amber) as a function of distribution s.d.. Distribution divergence (Δ_p.d.f._) is illustrated in black.

Here, I extend this framework by comparing the uniform noise assumption with Gaussian-distributed landmark uncertainty. Gaussian noise is a plausible and common choice, as it approximates the aggregate effect of multiple independent sources of variability – from peripheral transduction to central processing along the somatosensory pathway – consistent with the central limit theorem. To assess how much the shape of sensory noise (e.g., normal vs. uniform) affects model predictions, I computed the normalised average absolute difference in predicted performance between Gaussian and uniform noise models, matched for variance.

### The model shrinks input noise differences

In addition to Gaussian noise, I considered a rectangular (uniform) distribution as a physiologically and psychophysically grounded model of sensory uncertainty. This model assumes that timing uncertainty arises from a constant envelope amplitude threshold: a landmark (e.g., peak) is equally likely to be detected anywhere within the time window in which the envelope amplitude remains below that threshold. The width of this window maps into temporal variance, and the threshold itself is a free parameter of the model. This representation links naturally to envelope detection mechanisms in tactile processing and has been analytically derived in closed form [34]. It also approximates the effective temporal uncertainty when envelope discrimination threshold increases, particularly for sinusoidal envelopes, where the time-to-amplitude mapping flattens near the peak.

Despite its interpretability, the uniform distribution lacks some desirable properties of Gaussian noise. For instance, it cannot encode prior expectations about peak timing, and it lacks the statistical generality afforded by the central limit theorem. For these reasons, the Gaussian model is adopted for all behavioural fits. Nevertheless, it is important to assess the extent to which the model’s predictions depend on the assumed shape of sensory noise.

To isolate the model’s sensitivity to the input distribution, Figure 3A demonstrates predicted performance under uniform and Gaussian noise models, matched for variance, across a wide range of uncertainty levels (associated with amplitude thresholds from 0.1 to 0.9 relative to the peak amplitude). Figure 3B illustrates these comparison by plotting uniform-predicted performance (abscissa) against Gaussian predictions (ordinate), revealing that Gaussian predictions are consistently slightly higher, but the overall divergence remains modest. This difference was quantified using a normalised measure of prediction divergence (see Methods), as shown in Figure 3C. Across uncertainty levels, the mean performance difference was 16.7% ±2.3% (mean ± s.d.), substantially smaller than the total variation distance between the underlying noise distributions (39.5%), indicating that the model compresses large differences in input distribution shape into smaller differences in predicted behavioural output.

These results demonstrate that the tactile motion model is robust to the shape of the assumed sensory noise: while the rectangular model captures a distinct and meaningful physiological constraint, the predicted perceptual consequences differ only moderately from those of the Gaussian model.

### Asymmetry in psychometric functions: model predictions and empirical evidence

To further investigate model predictions of tactile motion perception, I examined whether the psychometric functions exhibit asymmetry in performance at phase offsets equidistant from the ambiguous conditions (0° and 180°). Specifically, we compared predicted performance at 30° versus 150° phase differences.

For the uniform noise model, the peak of the psychometric function was consistently shifted to values below 90°, reaching approximately 60° at high noise levels (e.g., *σ* = 0.6). In contrast, the Gaussian noise model produced a peak above 90° at low noise (up to ∼ 105°), which shifted to below 80° at higher noise regimes. This divergence in predicted peak locations demonstrates that the two models make qualitatively distinct predictions regarding phase-dependent performance (Figure 3D).

Although the behavioural paradigm did not permit direct localisation of the peak (since only 0° to 180° with 30° increments were tested), it enabled assessment of asymmetry by comparing performance at 30° and 150°. When quantifying asymmetry between 30° and 150°, the Gaussian model showed a maximum difference of Δ*P* = 8.3% (150° ¿ 30°) at *σ* = 0.14. A crossover point occurred at *σ* ≈ 0.385, where the predicted difference approached zero. For the uniform model, the asymmetry was much smaller, with a maximum Δ*P* = 4.16% at *σ* = 0.098 (amplitude threshold *δ* = 0.07), and a crossover at *σ* = 0.289 (threshold *δ* = 0.50). Thus, the Gaussian model not only produced larger asymmetry effects but also captured the reversal in relative performance across noise regimes. Figure 3E reveals systematic trend in the individual subject data consistent with the model predictions: performance at 150° exceeded that at 30° under conditions of lower overall performance, but the relationship reversed at higher performance levels. The empirical asymmetry was stronger in magnitude than that predicted by either model, with Gaussian predictions providing a closer match to the observed crossover pattern.

Together, these findings indicate that while both models predict phase-dependent asymmetry, the Gaussian model better captures the structure and magnitude of the empirical effects. Therefore, hereafter, the Gaussian formulation was adopted for subsequent analyses.

### Frequency-dependent temporal noise improves model fit

To evaluate the extent to which a single sensory timing noise parameter *σ* could account for performance across all frequencies (as compared to an individual fit per frequency as in Figure 4A), the model was fit to the performances across all conditions (frequencies and phase differences). The resulting fit were poor with an average root mean square error (r.m.s.e.) of 0.1156 (±0.0286 s.d.) across subjects. Consistently, when the model was fit to the cross-subject average performances, the r.m.s.e. was 0.3583, indicating that the model systematically failed to capture the frequency-dependent trends present in the data. Specifically, as shown in Figure 4B, the single-*σ* model underestimated performance at 1 and 1.5 Hz and overestimated it at 0.5 Hz. Note that r.m.s.e. is in the probability space, and and r.m.s.e. of 0.3583 means 35.8% error.

**Figure 4.**
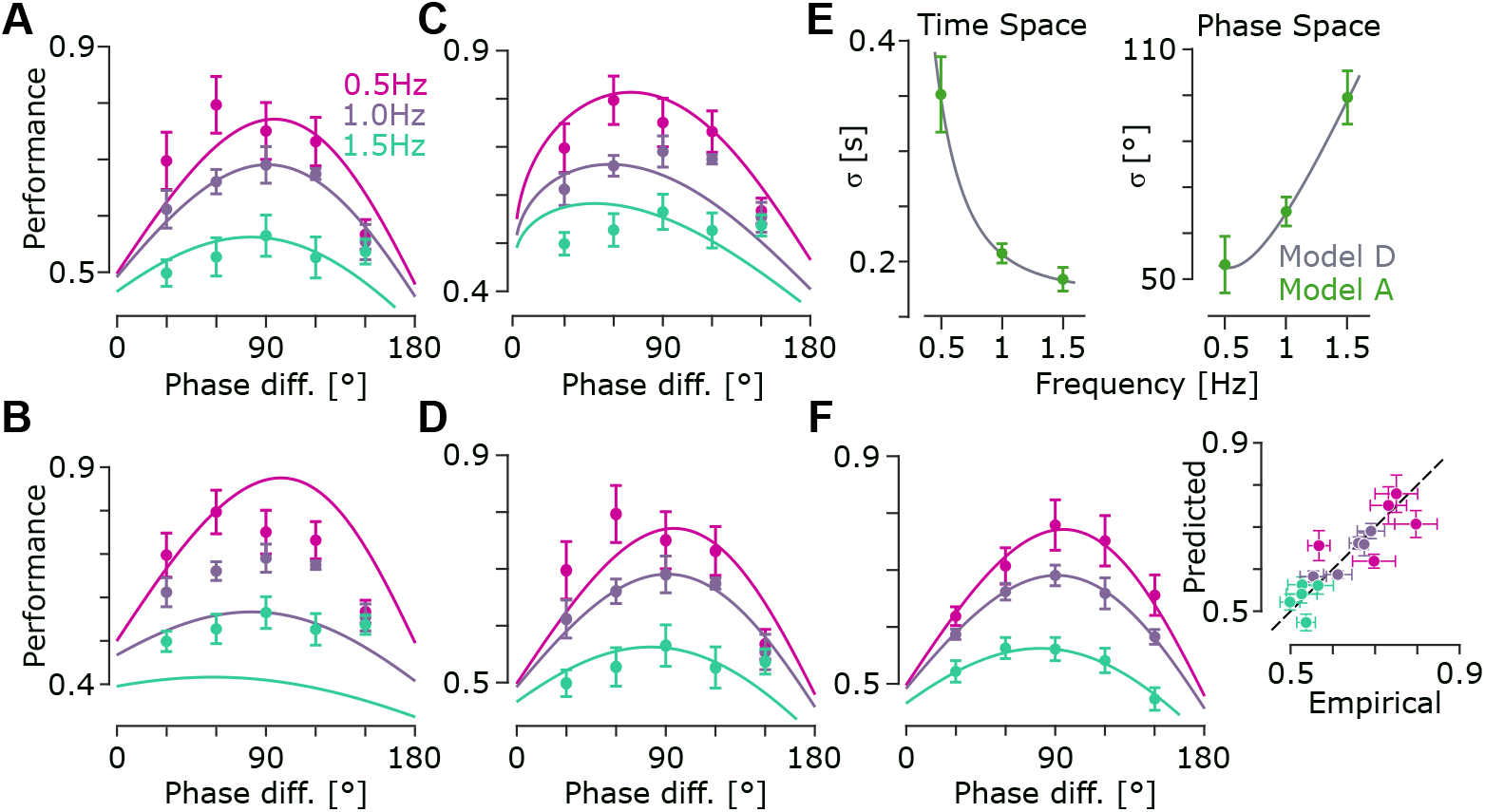
Model variants and frequency dependence of landmark timing uncertainty. **A**, Landmark model fitted independently to each modulation frequency. Markers represent empirical performance averaged across participants (*n* = 5); error bars indicate s.e.m. across participants. Solid curves show the best-fitting model predictions for each frequency. **B**, Same conventions as in **A**, but using a single timing uncertainty parameter (*σ*) fitted simultaneously across all modulation frequencies. **C**, Same conventions as in **A**, but fitting the landmark model with temporally correlated landmark uncertainty. **D**, Same conventions as in **A**, but fitting the dual uncertainty components model consisting of an amplitude-dependent and an amplitude-independent timing uncertainty. **E**, Frequency dependence of the fitted timing uncertainty in the time domain (left) and phase domain (right). Curves show predictions of the dual uncertainty components model. Green markers represent the independently fitted uncertainty estimates obtained from separate fits at each modulation frequency (**A**); error bars indicate s.e.m. across participants. **F**, Leave-one-phase-out generalisation analysis. For each phase difference, the model was fitted to all remaining phase differences across frequencies and used to predict the left-out condition. Solid curves show predictions from the dual uncertainty model (**D**). Markers represent the predicted performance for the omitted phase. Inset: predicted performance versus empirical performance at the left-out phase, averaged across participants. Error bars indicate s.e.m. across participants.

However, when the model was fit to individual frequencies separately (Figure 4A), allowing the model to capture frequency dependent nature of noise, the aggregate r.m.s.e. (computed as the square root of total squared error summed across frequencies and divided by the total number of conditions) on average were 0.0689 (±0.0175 s.d.), 40.4% lower than that of the single-*σ* model fit. Consistently, the net r.m.s.e. when fitting the model on the cross-subject average performances per frequency, was 0.0428, 88.0% lower than the corresponding single-*σ* model fit.

### Temporal noise correlations fail to capture structure and inflate noise to implausible levels

The inability of a single *σ* to fit performance across frequencies suggests the presence of temporal correlations in the timing uncertainties across landmarks. To test whether introducing temporal correlation improves model fit, I fitted an extended model in which timing noise across perceived landmark events decayed exponentially as a function of their temporal separation (see Methods). This model includes two free parameters: *σ*, the marginal standard deviation of timing noise, and *ρ*, a correlation coefficient that governs the strength of temporal dependency between peak times. Fitting the model to cross-subject average performance yielded a relatively good fit (r.m.s.e. of 0.0563), substantially outperforming the single-*σ* model (Figure 4C). The fitted parameters were *σ* = 5.92 s and *ρ* = 0.996, indicating strong positive temporal correlation and a high overall noise estimate. When fitted separately to each subject, the model produced average *σ* of 4.26 s (±2.02 s.e.m.) and *ρ* of 0.872 (±0.074 s.e.m.), with a mean r.m.s.e of 0.0827 (±0.0177 s.d.). Notably, the estimated *σ* values were substantially higher than the maximum estimated *σ* of 0.351 (±0.034 s.e.m.) across frequencies, predicting highly skewed phase-dependent patterns inconsistent with empirical data.

However, while the fits were quantitatively good, the inferred parameter values are not physiologically plausible. In particular, the fitted *σ* values are high relative to the modulation cycle durations (up to 2 s), and imply substantial timing variability. This reflects a compensatory behaviour in the model: to maintain low effective uncertainty over short intervals (e.g., 30° and 60°) across frequencies, the model compensates by inflating correlation across time, thereby spreading noise globally across the cycle. In effect, the model leverages its time-variable correlation structure to create tighter local estimates while keeping marginal variance high. This highlights that while the correlation-based model captures structured uncertainty, the resulting parameter values may not reflect a biologically realistic sensory noise profile.

### Dual internal uncertainty components

The frequency-dependent fits above (Figure 4A) suggest that temporal uncertainty is not constant across frequencies. Interestingly, while behavioural performance declines with frequency, the estimated temporal noise *σ* (fit independently for each frequency) decreases with frequency when expressed in absolute time units (Figure 4A). This pattern indicates that temporal precision improves with frequency, which may appear contradictory given the drop in behavioural accuracy.

To reconcile this, I propose a model with two additive sources of noise: one that decreases with frequency due to improved peak definition (e.g., amplitude-based detection noise [34]), and another that reflects a saturating central timing limit. Formally, total noise was modelled as the sum of a power-law term and a constant term:

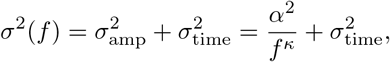

where *σ*(*f* ) is the net timing uncertainty, *f* is modulation frequency, *α* controls amplitude-dependent uncertainty, *σ*_amp_ represents amplitude-based detection noise and *σ*_time_ represents the central resolution limit.

Fitting this dual-noise model yielded remarkably close estimates to those obtained from per-frequency fits – group-level r.m.s.e. of 0.0428, with summed squared error (s.s.e.) of 1.02 × 10^—10^ between the dual-noise model prediction and per-frequency fit (see Figure 4A vs. D). The group-level estimate of *σ*(*f* ) matched the average of per-frequency *σ* values across subjects (s.s.e. of 1.31 × 10^—5^). Across subjects, the dual-noise model fit the data with an average r.m.s.e. of 0.0689 (± 0.0175 s.d.) and group-level r.m.s.e. of 0.0428 for fit to cross-subject average data.

Figure 4E shows the estimated *σ*(*f* ) as a function of frequency in both time and phase units. In time units, *σ*(*f* ) decreases with frequency, matching the per-frequency fits. In contrast, when transformed to phase units, the uncertainty increases (approximately) linearly with frequency. This dual trend is consistent with a two-component noise structure: a fast-decaying amplitude-related component and a fixed temporal resolution limit. The power-law exponent *κ* fitted to group data was 2.15, with a cross-subject average of 2.54 ± 0.41 (s.e.m.). The linear phase-domain slope was 0.16 ± 0.02 s.e.m. s per Hz (group level: 0.17 s per Hz), or equivalently, 58.9°±6.7° per Hz, matching the behavioural performance decline.

### Model generalises accurately to unseen phase differences

To assess the generalisability of the model, I tested its ability to predict performance at unseen phase differences. Specifically, for each phase difference, the model was fit to all other phase values across all frequencies, and then used to predict performance at the held-out phase. This process was repeated for each phase value, and predictions were averaged across subjects (see Methods).

As shown in Figure 4F, the model’s predictions closely matched those obtained when fitting to the full dataset. The inset panel compares the predicted performance at held-out phase values with the actual empirical data, averaged across subjects. Remarkably, the predicted values aligned closely with both the empirical measurements and the full-data model fits, demonstrating that the model captures systematic structure across phase and frequency, and generalises well to unseen conditions.

### Decreased Internal Uncertainty with Sharply Defined Landmarks

To test the prediction that sharper temporal landmarks reduce internal timing uncertainty, a separate temporal order judgement (TOJ) experiment was conducted. In a 2AFC task, participants judged whether the perceived peak of a vibrotactile stimulus occurred before or after a brief auditory probe (10-ms auditory tone) presented at temporal offsets of ± 50, ± 150 or ± 300 ms relative to the vibrotactile landmark (here, the envelope peak). Vibrotactile stimuli consisted of a single modulation cycle (2 s duration; 0.5 Hz) with either a broad sinusoidal envelope or a sharper envelope defined by a sinusoid raised to the fifth power. (Figure 5A).

**Figure 5.**
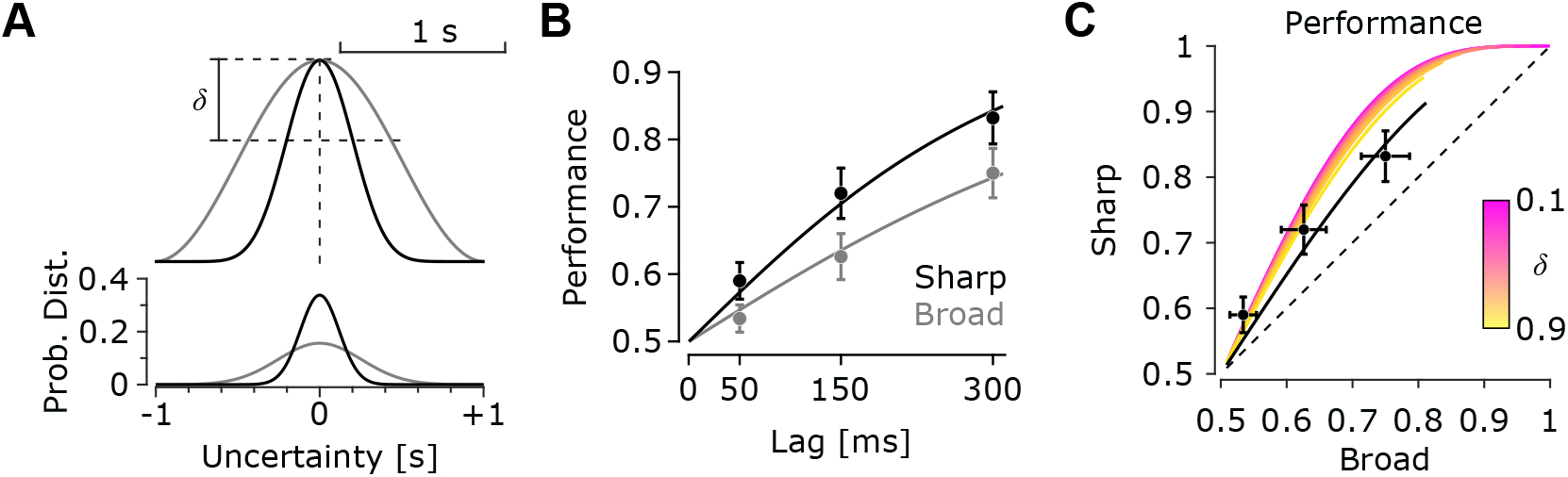
Sharper temporal landmarks reduce perceptual uncertainty and support a dual uncertainty components account. **A**, Top: broad (grey) and sharp (black) envelopes in the temporal order judgement (TOJ) experiment. The dashed lines indicate an example amplitude threshold *δ*, relative to the envelope peak. The threshold defines a temporal uncertainty window surrounding the peak, with its width depends on envelope sharpness. Bottom: equivalent Gaussian probability density functions describing landmark timing uncertainty for the two envelopes. The Gaussian distributions were obtained by matching the variance of a uniform distribution defined over the corresponding threshold-derived temporal windows. **B**, Temporal order discrimination performance as a function of the temporal separation between the auditory probe and tactile landmark. Markers represent participant means; error bars indicate s.e.m.. Curves show participant-specific conditional predictions from the hierarchical generalised linear mixed-effects model, averaged across participants. **C**, Temporal order discrimination for the sharp envelope plotted against the corresponding performance for the broad envelope. Markers represent participant means; error bars indicate s.e.m.. Coloured curves show landmark model predictions for amplitude threshold *δ* from 0.1 to 0.9 (increments of 0.1), where landmark timing uncertainty is derived analytically from the threshold-defined temporal uncertainty window surrounding the envelope peak. The black curve corresponds to the dual uncertainty components model, incorporating both the amplitude-dependent landmark uncertainty and the shared amplitude-independent uncertainty component.

**Figure 6.**
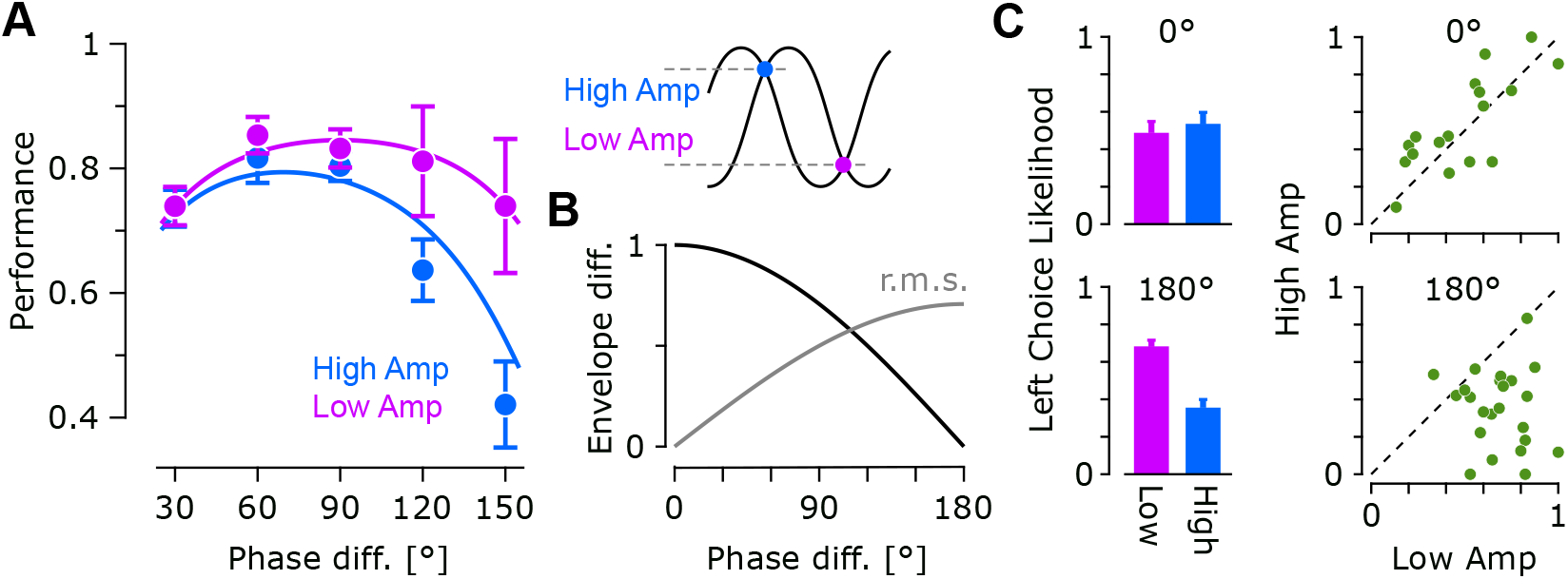
Phase-dependent effects of the initial envelope state on tactile motion perception. **A**, Motion discrimination performance for the two iso-amplitude onset conditions as a function of phase difference. Symbols represent participant means; error bars indicate s.e.m.. Curves show population-level predictions of the hierarchical generalised linear mixed-effects model obtained by integrating over the estimated subject-level random-effect distribution using Monte Carlo integration (10,000 samples). Inset: representative stimulus envelopes illustrating the two iso-amplitude onset conditions. **B**, Initial amplitude difference (black) and root-mean-square (r.m.s.) of instantaneous difference (grey) between the two vibration envelopes as a function of phase difference. The initial amplitude difference corresponds to the absolute difference between envelope amplitudes at stimulus onset, whereas the r.m.s. curve quantifies the average instantaneous difference over one modulation cycle (2 s). Note that the initial amplitude difference is largest at small phase differences and approaches zero at 180°, opposite to the behavioural modulation observed in **A. C**, Choice preference for the ambiguous phase conditions. Left panels show the proportion of leftward responses (mean ± s.e.m.) for the two initial conditions at 0° (top) and 180° (bottom). Right panels show the corresponding participant-level data; each marker represents one participant. At 180°, the two initial conditions correspond to identical envelope amplitudes and differ only in the direction of the envelope change (time derivative) during the first 500 ms preceding the first landmark peak preceding the first landmark peak.

Temporal order discrimination improved with increasing temporal separation between the auditory probe and the tactile landmark, with the sharper envelope exhibiting consistently higher accuracy (Figure 5B). For the broad envelope (sine envelope exponent of 1), mean performance increased from 53.4% ±2.0% s.e.m. at 50 ms to 62.6% ± 3.4% at 150 ms and 75.0% ±3.7% at 300 ms. For the sharper envelope (sine exponent of 5), performance was consistently higher, increasing from 59.0% ±2.7% at 50 ms to 72.0% ±3.7% at 150 ms and 83.2% ±3.9% at 300 ms. These findings were confirmed by a hierarchical trial-level analysis of accuracy using a binomial generalised linear mixed-effects model with a logit link and subject-specific random slopes. Discrimination accuracy increased significantly with temporal separation from the tactile peak (*β* = 3.76 ± 0.66 s.e.m., 95% CI [2.47, 5.05], *p <* 10^—7^). Critically, the effect of temporal separation was significantly stronger for the sharper envelope (interaction term: *β* = 2.12 ± 0.45, 95% CI [1.23, 3.02], *p <* 10^—5^), indicating improved temporal precision when the tactile peak was more sharply defined.

I further quantified landmark timing uncertainty using a hierarchical generalised linear model of response likelihood as a function of temporal offset, assuming a binomial distribution and probit link. Under the Gaussian uncertainty assumption of the landmark model, the probit slope provides a direct estimate of the landmark timing uncertainty parameter, *σ*. Sensitivity was significantly higher for the sharper envelope (Δ*β* = 1.19 ± 0.26, 95% CI [0.68, 1.71], *p <* 10^−5^). The corresponding uncertainty estimates were 452 ± 36 ms for the broad envelope and 294 ± 17 ms for the sharper envelope, representing a 35.0% reduction in landmark timing uncertainty.

The uncertainty estimate obtained in this TOJ task includes contributions from both tactile and auditory timing as well as temporal comparison processes. However, because the auditory probe was a brief 10-ms tone with a sharply defined onset, its temporal uncertainty is likely substantially smaller than that of the tactile landmark with 100 Hz carrier (10 ms per cycle) modulated by a slowly varying envelope. Consequently, the uncertainty estimate from TOJ task primarily reflects tactile landmark uncertainty and may be viewed as an upper bound on the tactile uncertainty parameter. Notably, in this context, despite the different participant cohorts and task requirements, the uncertainty estimate obtained from the TOJ task (452 ± 36 ms) was of comparable magnitude to the value independently estimated from the motion-discrimination model at 0.5 Hz (351 ± 34 ms), consistent with both tasks probe a common underlying process of tactile landmark timing uncertainty.

Because the auditory reference stimulus was identical across conditions, the improved performance for the sharper envelope is most parsimoniously explained by greater precision in estimating the timing of the tactile landmark itself. The model-based estimates of uncertainty demonstrate that sharpening the envelope substantially reduces landmark timing uncertainty, providing direct experimental support for the landmark-based motion perception framework. Importantly, the estimated uncertainty for the broad sinusoidal envelope corresponds to the identical vibrotactile stimulation pattern used in the motion-perception experiments, linking the timing precision measured here to the uncertainty parameter governing tactile motion discrimination.

### Dual Uncertainty Components of Landmark Timing

To examine the extent to which the landmark account based on envelope geometry quantitavily explains the observed improvement in temporal precision, I used the landmark framework to predict temporal precision from the temporal window surrounding the envelope peak. Specifically, for a given amplitude threshold *δ*, the interval over which the envelope falls within the threshold margin of its peak was converted to an equivalent Gaussian uncertainty by matching the variance of the corresponding uniform distribution. Across a broad range of threshold values, the model correctly predicted lower uncertainty for the sharper envelope than for the broad envelope. However, the single-parameter model based on envelope threshold-dependent uncertainty alone consistently overestimated the separation between the two conditions (Figure 5C), indicating that although the model captured the direction of the effect, it did not fully account for the compressed difference observed in the behavioural data between the two envelopes.

We therefore considered a model with two uncertainty components. The first component was an amplitude-dependent uncertainty term, *σ*_amp_, determined by the temporal uncertainty about the envelope peak, derived directly from the envelope geometry within the landmark framework. The second component was an amplitude-independent uncertainty term, *σ*_0_, shared across envelope conditions. Under this formulation,

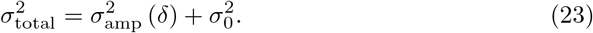

Using the uncertainty estimates obtained independently from the hierarchical probit psychometric analysis (*σ*_broad_ = 452 ± 36 ms and *σ*_sharp_ = 294 ± 17 ms), the threshold parameter and shared uncertainty component were solved simultaneously. This yielded a threshold value of *δ* = 0.775 and a shared uncertainty component of *σ*_0_ = 219 ms. The resulting model reproduced the observed psychometric performance with high accuracy (r.m.s.e. of 0.016), substantially improving the agreement between theory and empirical data compared to the amplitude-dependent component alone. These findings indicate that the temporal precision measured in the TOJ task is not determined solely by landmark sharpness, but reflects the combined influence of an amplitude-dependent component corresponding to the landmark extraction based on the stimulus geometry and an amplitude-independent component shared across conditions.

Together, these findings provide direct experimental support for the landmark-based framework and the proposed dual uncertainty components account. Sharper envelope peaks reduce the amplitude-dependent component of uncertainty associated with landmark extraction, while the shared uncertainty component captures additional sources of temporal imprecision common to both envelope conditions, including potential contributions from auditory timing, cross-modal comparison, and decision-related uncertainty. The agreement between the independently estimated probit uncertainties and the two-component landmark model further links the temporal precision measured in the TOJ task to the uncertainty parameter governing tactile motion discrimination.

### Initial Envelope State Reveals Distinct Sources of Motion Ambiguity and Supports Landmark-Based Decisions

To characterise the extent to which perceptual decisions were influenced by the initial envelope state versus by the subsequent sequence of envelope landmarks, I separated trials according to the two iso-amplitude onset conditions (see Methods). At these two onset conditions, the two vibrations started at identical amplitudes (Figure 6A inset), but differed in the subsequent temporal sequence of envelope changes and peaks. The two onset conditions are referred to as the high and low initial-amplitude conditions, following the nomenclature introduced in the Methods. At 180°, the same labels were used for consistency, although the two onset conditions differ only in the temporal sequence of envelope changes and landmark peaks rather than in their initial amplitudes.

The influence of the initial envelope state depended strongly on phase difference (Figure 6A). At smaller phase offsets (30°–90°), discrimination performance was nearly identical between the two onset conditions, with mean differences (high minus low) of -0.3±2.2%, -3.7±2.8% and -2.8±2.3% at 30°, 60° and 90°, respectively. In contrast, progressively larger differences emerged at higher phase offsets (¿90°). At 120°, performance was 63.7±4.9% for the high-amplitude initial-state condition and 8.8% for the low-amplitude initial-state condition, corresponding to a difference of -17.5±11.6%. At 150°, performance was 42.1±6.9% and 74.0±10.8%, corresponding to a reduction of 31.9±17.4%. Thus, the influence of the initial envelope state was negligible over the temporal-order regime (¡90°) but increased substantially as phase difference approached the region where cross-cycle correspondence becomes increasingly ambiguous (Figure 6A).

These observations were confirmed using a hierarchical trial-level generalised linear mixed-effects model with a binomial response distribution and subject-specific random slopes. The model included a phase-dependent motion evidence sin (*ϕ*) together with a phase-dependent modulation associated with the initial stimulus state and was compared with a nested model containing only the phase-dependent motion cue. Incorporating the initial-state modulation significantly improved the model fit (likelihood-ratio test: *χ*^2^(1) = 28.96, *p <* 10^—7^; Δ*AIC* = 27; Δ*BIC* = 20). The phase-dependent motion cue was strongly associated with performance (*β* = 2.33 ± 0.28, 95% CI [1.79, 2.87], *p <* 10^—16^), indicating that performance increased systematically with phase evidence. Importantly, the interaction between the phase evidence and the initial envelope state was significantly negative (*β* = −1.12 ± 0.14, 95% CI [-1.40, -0.85]; *p <* 10^—14^), demonstrating that the detrimental modulation effect of the high initial-state condition increased progressively with phase difference. Population-level predictions obtained by integrating over the estimated subject-level random-effects distribution closely reproduced the observed behavioural trend of minimal modulation at small phase differences and progressively larger effects at higher phase differences (Figure 6A). The estimated dispersion parameter of 0.999 indicated no evidence of overdispersion relative to the binomial assumption (*n* = 7,105 trials).

The phase-dependent modulation produced by the initial envelope state closely matched the transition predicted by the landmark model between the two sources of perceptual ambiguity. At smaller phase offsets, discrimination is primarily limited by temporal-order uncertainty between neighbouring landmarks and is largely unaffected by the initial stimulus state. As phase difference increases beyond 90°, temporal-order judgements become progressively more reliable, whereas assignment of landmarks to the correct modulation cycle becomes increasingly ambiguous. The growing behavioural influence of the initial envelope state therefore emerges specifically within the regime where cross-cycle correspondence dominates perceptual uncertainty, consistent with the two computational components of the landmark framework.

Although the two onset conditions differed only in the initial state of the envelopes, their perceptual effects were opposite to those expected from a computation based solely on the initial direction of envelope change or instantaneous amplitude differences. Instead, the modulation followed the temporal organisation of the subsequent landmark sequence. This iwas further supported by the opposite dependence of perceptual modulation (Figure 6A) and instantaneous envelope difference (Figure 6B), indicating that perceptual decisions are determined primarily by the temporal sequence and correspondence of landmark events rather than by instantaneous envelope amplitude or motion energy.

### Initial Envelope State Biases Perceptual Choices at 180° but not at 0°

The geometry of the stimulus at 0° and 180° phase differences provides an informative test of how the initial stimulus state influences perceptual decisions. At both phase differences, motion discrimination accuracy is undefined because no objectively correct direction exists. Consequently, perceptual responses were quantified as the proportion of leftward choices. Although the two onset conditions were defined using the same iso-amplitude criterion as for all other phase differences, they differ qualitatively in these two conditions. At 0°, the onset conditions correspond to the largest possible difference in initial envelope amplitude while remaining identical in phase throughout the trial. In contrast, at 180°, the two onset conditions have identical envelope amplitudes and differ only in the subsequent sequence of envelope changes and landmark peaks. These conditions, therefore, dissociate the influence of instantaneous envelope amplitude and its derivative from that of the temporal sequence of landmark events.

At 0°, the proportion of leftward responses was 53.5±6.1% for the high-amplitude initial-state condition and 48.8±6.1% for the low-amplitude initial-state condition, indicating little difference in perceptual choices between the two conditions (Figure 6C). A hierarchical trial-level generalised linear mixed-effects model with subject-specific random intercepts confirmed that the initial stimulus state did not significantly influence responses (*β* = 0.24 ± 0.19, 95% CI [-0.13, 0.61], *t*(507) = 1.29, *p* = 0.196). The intercept was also not significantly different from zero (*β* = −0.10 ± 0.25, 95% CI [-05·9, 0.68], *t*(507) = −0.41, *p* = 0.680), indicating that choices were not systematically biased away from chance. The estimated dispersion parameter of 0.958 indicated no evidence of over-dispersion relative to the binomial assumption (*n* = 509) trials.

In contrast, a significant effect emerged at 180° (Figure 6C). The proportion of left responses was 35.5±4.3% for the onset condition labelled *high* and 68.2±3.3% for the onset condition labelled *low*, corresponding to a difference of 32.7 percentage points. Although these conditions are referred to as high and low for consistency with the remaining phase differences, they have identical initial envelope amplitudes and differ only in the subsequent temporal sequence of envelope changes and landmark peaks. The observed bias was consistent with the temporal sequence of landmark peaks but opposite to the direction predicted by the initial envelope change during the first 500 ms. A hierarchical generalised linear mixed-effects model revealed a strong effect of the initial envelope state (*β* = −1.42 ± 0.16, 95% CI [-1.73, -1.12], *t*(735) = −9.09, *p <* 10^—18^). The negative coefficient indicates that the high onset condition substantially reduced the probability of reporting leftward motion relative to the low onset condition. The intercept was significantly positive (*β* = 0.80 ± 0.12, 95% CI [0.56, 1.04], *t*(735) = 6.53, *p <* 10^—9^), indicating a significant leftward response bias in the reference (low onset) condition. The estimated dispersion parameter was 0.988, indicating no evidence of over-dispersion relative to the binomial assumption (*n* = 737 trials).

The two ambiguous phase conditions (0° and 180°) therefore exhibited qualitatively different behaviour. At 0°, the onset condition had no systematic influence on perceptual choices despite producing the largest possible difference in initial envelope amplitude. In contrast, at 180°, where the two onset conditions differed only in the subsequent sequence of envelope changes and landmark peaks, the onset condition produced a pronounced choice bias. This bias followed the temporal sequence of landmark peaks rather than the direction implied by the initial envelope change during the first 500 ms before the first peak. The observed selective bias, along with the opposite dependence of behavioural modulation of performance (Figure 6A) and instantaneous envelope difference (Figure 6B), argue against a computation based solely on instantaneous envelope amplitude or motion energy and instead support a computation based on the temporal sequence and correspondence of landmark events.

## Discussion

The present study investigated the computational structure of temporal uncertainty underlying phase-based tactile motion perception. Building on the previously established landmark framework [34, 35], the findings show that perceptual performance is governed by multiple sources of uncertainty rather than by a single internal timing noise process. Behavioural measures of accuracy, response time, confidence, and confidence entropy exhibited systematic signatures of perceptual uncertainty across phase differences. Computational modelling further demonstrated that a single uncertainty component cannot fully account for the observed behavioural structure. Instead, the data are best explained by two complementary sources of timing uncertainty: an amplitude-dependent component associated with extracting temporal landmarks from the vibration envelope, and an amplitude-independent component that limits temporal precision across stimulus conditions. An independent temporal order judgement experiment directly validated these uncertainty estimates, while manipulation of the initial stimulus state demonstrated that perceptual decisions follow the temporal sequence of landmark events rather than instantaneous envelope amplitude or motion energy.

These findings strengthen the view that tactile motion perception is fundamentally a problem of temporal inference. Rather than relying on continuous estimates of instantaneous stimulus energy, the nervous system appears to extract salient temporal landmarks, establish their correspondence across digits, and infer motion direction from their relative timing. Central to this computation is the ability to detect and compare temporally structured vibration signals, a strategy that has analogues across sensory systems and species. For example, arachnids localise prey from complex substrate vibrations transmitted through webs and other mechanical media using highly specialised vibration-sensitive mechanoreceptors [43]. Likewise, vertebrate tactile systems contain specialised mechanoreceptors for encoding vibrotactile signals, with Meissner’s and Pacinian corpuscles providing complementary representations of dynamic skin deformation and high-frequency vibration [44–48]. In particular, Pacinian afferents have been implicated in vibrotactile frequency and pitch perception in both rodents and humans [32, 33, 49]. Consistent with these observations, previous work from our laboratory demonstrated that rodents accurately discriminate vibration amplitude and frequency using their whiskers and that these stimulus features are represented with high temporal precision in primary somatosensory cortex [50–52]. The present findings extend these principles by showing that the relative timing of vibrotactile envelope landmarks, rather than instantaneous vibration amplitude itself, provides a robust computational substrate for tactile motion perception.

### Landmark Timing and Correspondence Underlies Tactile Motion Perception

The present findings provide converging behavioural and computational evidence that tactile motion perception in this paradigm is governed primarily by the temporal organisation of salient envelope landmarks rather than by instantaneous envelope amplitude or continuously varying motion energy. The strongest support for this conclusion comes from the initial-state manipulation (Figure 6). Although the two onset conditions start at identical envelope amplitudes, they differed in the subsequent temporal sequence of envelope changes and landmark peaks. At phase differences*>*90°, as the inter-peak intervals increase progressively with phase offset, the direction implied by the initial envelope change was opposite to that defined by the subsequent landmark sequence. Nevertheless, perceptual choices consistently followed the landmark-defined interpretation. This perceptual influence of the initial state increased specifically within the phase range where the landmark model predicts increasing correspondence ambiguity, while remaining negligible in the temporal-order regime (*<*90°). These findings, argue against a computation based solely on instantaneous envelope amplitude, correlation or motion energy, and instead support one based on the temporal ordering and correspondence of salient landmark events.

landmark-based computation belongs to a broader class of feature-based perceptual mechanisms in which motion and temporal judgements can depend on the detection, tracking, or matching of salient events or features rather than on continuous readout of raw stimulus intensity [26, 53–56]. Similar principles are well established in auditory perception, where stimulus onsets and other transient temporal cues strongly influence sound localisation, precedence effects, temporal order judgements, and the segmentation of acoustic events [57–63]. In visual motion perception, feature-based and correspondence-based mechanisms provide an alternative to purely luminance-energy based motion computation, with motion sometimes inferred by matching salient features, object tokens, or marked locations across time [53–56, 64]. In tactile perception, apparent motion similarly depends on the temporal relationship between discrete stimulation events, including inter-stimulus onset interval, stimulus duration, and the spatial sequence of discrete tactile events [15, 17, 30]. The present findings extend these principles to continuous vibrotactile stimulation by suggesting that the nervous system extracts salient temporal landmarks from each vibration envelope and infers motion from their relative timing and correspondence rather than from instantaneous envelope amplitude differences [34, 35].

### Temporal Order and Landmark Correspondence Represent Distinct Sources of Perceptual Ambiguity

The landmark model separates tactile motion inference into two probabilistic computations: first, estimating the temporal order of corresponding landmarks, and, second, assigning those landmarks to the appropriate modulation cycle [34, 35]. This distinction is consistent with classical work on temporal order judgement, which demonstrated that judging the order of brief sensory events is fundamentally limited by temporal resolution, as well as later work showing that temporal-order processing depends on stimulus characteristics such as frequency and waveform [57, 60, 65]. Consistent with these findings, motion discrimination in the present study exhibited a strong dependence on modulation frequency (Figure 4), which can be accounted for by an amplitude-dependent component of landmark timing uncertainty. This interpretation is independently supported by the TOJ experiment, in which sharper envelope waveforms reduced landmark timing uncertainty (Figure 5), and is further consistent with the enhanced motion discrimination previously observed for exponential compared with sinusoidal envelope modulation [35].

In the present task, the dominant source of perceptual uncertainty is temporal-order estimation when phase differences are smaller than 90°, because corresponding landmarks occur close together in time and their perceived order is therefore unreliable. By contrast, as the phase difference approaches 180°, temporal order becomes progressively more reliable, whereas the dominant source of uncertainty shifts towards establishing correspondence between landmarks across neighbouring modulation cycles, because a landmark from one vibration can be assigned with similar probability to either the preceding or following cycle of the other vibration [<u>34, 35</u>]. This transition is reflected in the asymmetric psychometric functions (Figure 3), the distinct profiles of confidence (Figure 2C) and selective choice bias (Figure 6C) observed at 180°. At 0°, ambiguity arises because no directional temporal order exists between corresponding landmarks. At 180°, ambiguity arises despite strong temporal structure because landmark correspondence becomes uncertain. Together, these findings demonstrate that similar behavioural ambiguity may arise from fundamentally different computational sources depending on the phase relationship between the vibration envelopes. The transition from temporal-order uncertainty to correspondence ambiguity provides a unified framework for the characteristic asymmetry of the psychometric function and the distinct behavioural signatures observed across phase differences.

The mathematical formulation of the landmark model naturally explains this transition. The decision variable is not determined by temporal order alone, but by the joint probability that landmarks are both correctly ordered and correctly assigned to the same modulation cycle (Eq. **??**). The temporal-order term quantifies the probability that a landmark at one fingertip is perceived before the corresponding landmark at the other fingertip, whereas the correspondence term quantifies the probability that the relevant landmarks are assigned to the same cycle rather than neighbouring cycles. Their joint probability therefore defines the reliability of a directional interpretation for each landmark sequence. In this sense, the model formalises tactile motion perception as probabilistic inference over uncertain temporal events, consistent with broader Bayesian accounts in which perceptual decisions depend on representing and combining uncertainty in sensory evidence [1–3].

### Independent Validation of Landmark Timing Uncertainty

The previous landmark model treated landmark timing uncertainty as a latent computational parameter inferred from motion discrimination performance [34]. The present study demonstrates that this parameter corresponds to a measurable psychophysical quantity that can be quantified independently of the motion task. Specifically, the TOJ experiment showed that sharper envelope peak improved temporal discrimination and reduced the estimated landmark timing uncertainty by approximately 35%. Additionally, the uncertainty estimates obtained from the independent TOJ experiment matched those estimated from the motion discrimination task despite the two experiments differing in behavioural demands, participant cohorts, and decision processes. This agreement indicates that the uncertainty parameter governing the landmark model reflects a genuine property of tactile temporal processing rather than merely serving as a free fitting parameter.

The TOJ experiment also demonstrated that landmark timing uncertainty depends systematically on the temporal structure of the stimulus. Sharper envelope waveforms produced more precise temporal localisation of landmarks, consistent with classical studies showing that temporal order judgements depend not only on central temporal resolution but also on stimulus characteristics such as waveform, duration, and temporal salience [57, 60, 65]. Similar stimulus-dependent changes in temporal precision have been reported across auditory and tactile perception, where sharper temporal transients produce more reliable perceptual timing than slowly varying stimuli [63, 66]. The present findings therefore extend these observations by linking stimulus-dependent temporal precision directly to the computational parameter governing tactile motion perception. Rather than being inferred solely from motion perception, landmark timing uncertainty can be independently measured, manipulated through stimulus geometry, and quantitatively related to perceptual performance across distinct behavioural tasks. This provides a strong validation for a computational model of sensory decision-making, in which the principal model parameter corresponds directly to an independently measurable psychophysical quantity.

### Dual Components of Landmark Timing Uncertainty

A single uncertainty component derived from envelope geometry through a single parameter [34] consistently captured the qualitative direction of behavioural effects but systematically overestimated their magnitude. This pattern emerged independently in three experiments. First, frequency-dependent modelling of motion discrimination showed that uncertainty predicted solely from landmark geometry produced changes larger than those observed behaviourally (Figure 4B). Second, the temporal order judgement experiment demonstrated that waveform-dependent changes in landmark sharpness produced a compressed behavioural difference relative to that predicted by a single amplitude-dependent uncertainty component. Third, the previously reported comparison between exponential and sinusoidal envelope modulation showed the same qualitative pattern, with the original single-parameter landmark model systematically overestimating the behavioural improvement associated with sharper envelopes [35]. The convergence of these independent observations strongly suggests that landmark timing uncertainty cannot be explained by stimulus geometry alone.

The present results indicate that landmark timing uncertainty is better described by two additive components. The first depends directly on stimulus geometry and reflects the uncertainty associated with extracting temporal landmarks from the vibration envelope. This component decreases as envelope peaks become sharper and therefore predicts improved temporal precision for higher modulation frequencies and sharper waveforms. The second component is independent of envelope shape and represents a residual source of temporal uncertainty common to all conditions. This shared component limits the achievable temporal precision even when landmarks become increasingly well defined.

These two components produce opposite trends when uncertainty is expressed in the time and phase domains (Figure 4E). In absolute time, the amplitude-dependent component decreases as modulation frequency increases because landmarks become temporally sharper. However, when expressed in phase coordinates, uncertainty increases because the same absolute timing error occupies a progressively larger fraction of the modulation cycle. The resulting interaction naturally explains the observed frequency dependence of tactile motion perception while remaining consistent with the independent TOJ measurements (Figure 5C).

The existence of multiple uncertainty sources is also consistent with broader psychophysical evidence suggesting that perceptual timing reflects both stimulus-dependent and stimulus-independent limitations. Fink et al. demonstrated that temporal order judgements contain separable components associated with stimulus-specific processing and central timing mechanisms [60]. Likewise, Bayesian theories of perception propose that sensory uncertainty arises from multiple stages of neural processing, each contributing to the reliability of the final perceptual estimate [1–3]. The present results provide direct computational evidence for this principle in tactile motion perception by independently identifying, quantifying, and validating two distinct components of landmark timing uncertainty.

### Landmark Correspondence as the Unit of Evidence for Perceptual Decisions

The considerable variability in response times observed both within and across participants (Figure 1), together with the systematic relationships between response time, confidence, confidence entropy, and discrimination accuracy (Figure 2),suggest that tactile motion perception is unlikely to arise from a fixed-latency computation. Rather, these behavioural signatures are consistent with sequential evidence accumulation models, in which sensory evidence is integrated over time until a decision criterion is reached [ . While the present experiments were not designed to fit diffusion-type decision models explicitly, they provide insight on the form of the sensory evidence that such models would accumulate.

In the landmark framework, the fundamental unit of sensory evidence is the successful correspondence of a landmark pair across the two fingertips. The reliability of each correspondence is determined jointly by the probability of correct temporal ordering and the probability that the two landmarks are assigned to the same modulation cycle (Eq. 12 and Eq. 13, respectively). Their joint probability therefore quantifies the reliability of a directional interpretation associated with each landmark sequence (Eq. 18). Motion evidence accumulates as successive landmark correspondences become available throughout stimulation, with each contribution weighted according to its temporal reliability and correspondence uncertainty. This formulation links the decision variable to the probabilistic computations developed in the landmark model rather than introducing an additional decision-stage parameterisation.

This framework also generates experimentally testable predictions that distinguish landmark-based and correlation-based motion computations. Correlation-based accounts assume that evidence evolves continuously as temporal correlations between the two vibration signals are integrated over time, leading to the unimodal reaction-time distributions characteristic of diffusion-like decision processes [**?, ?**]. In contrast, the landmark framework predicts that evidence becomes available only when informative landmark correspondences are detected. If a single landmark dominates each modulation cycle, response-time distributions should exhibit modes aligned with successive landmark events. More generally, when multiple informative landmarks contribute within a cycle, each landmark correspondence contributes an additional burst of sensory evidence, with the number and temporal locations of response-time modes reflecting the underlying landmark structure. Characterising response-time distributions under conditions of systematically manipulated landmark correspondence therefore provides a direct experimental approach for distinguishing between continuous correlation-based computations and discrete landmark-based evidence accumulation. Although the present dataset contains insufficient trials per participant to resolve such distributions reliably, the phase-dependent influence of the initial state provides behavioural evidence favouring landmark-based computations over instantaneous correlation or energy-based accounts.

## Conclusion

The present study demonstrates that tactile motion perception can be understood as probabilistic inference over uncertain temporal landmarks. By combining behavioural experiments, computational modelling, and an independent temporal order judgement task, the present study showed that the uncertainty governing tactile motion perception is neither fixed nor unitary. Instead, landmark timing uncertainty comprises multiple computational components that arise from both stimulus-dependent landmark extraction and stimulus-independent temporal processing. These components jointly account for the frequency dependence of motion perception, the effects of waveform sharpness, the characteristic asymmetry of the psychometric function, and the distinct computational roles of temporal-order estimation and cross-cycle landmark correspondence. Importantly, the principal uncertainty parameter of the landmark model was independently validated using a separate TOJ paradigm, providing direct experimental support for its physiological interpretation rather than treating it as a purely descriptive fitting parameter.

Broadly, the landmark framework establishes phase-based tactile motion perception as a tractable model system for investigating the computational structure of perceptual uncertainty. Unlike many perceptual paradigms in which different sources of internal variability are difficult to separate experimentally, the present approach allows stimulus geometry, temporal landmark reliability, correspondence ambiguity, and decision-related uncertainty to be manipulated and quantified independently. These principles are likely to extend beyond tactile motion perception to other sensory systems in which perception depends on the temporal correspondence of discrete events. By linking computational theory with independently measurable psychophysical quantities, the present work provides a framework for studying how structured sensory uncertainty gives rise to perceptual decisions within and beyond the tactile domain.

## References

1. Knill DC, Pouget A. The Bayesian brain: the role of uncertainty in neural coding and computation. TRENDS in Neurosciences. 2004;27(12):712–719.

2. Ma WJ, Beck JM, Latham PE, Pouget A. Bayesian inference with probabilistic population codes. Nature neuroscience. 2006;9(11):1432–1438.

3. Pouget A, Beck JM, Ma WJ, Latham PE. Probabilistic brains: knowns and unknowns. Nature neuroscience. 2013;16(9):1170–1178.

4. Ma WJ. Signal detection theory, uncertainty, and Poisson-like population codes. Vision research. 2010;50(22):2308–2319.

5. Moreno-Bote R, Beck J, Kanitscheider I, Pitkow X, Latham P, Pouget A. Information-limiting correlations. Nature neuroscience. 2014;17(10):1410–1417.

6. Aitchison L, Lengyel M. The Hamiltonian brain: Efficient probabilistic inference with excitatory-inhibitory neural circuit dynamics. PLoS computational biology. 2016;12(12):e1005186.

7. Ni AM, Ruff DA, Alberts JJ, Symmonds J, Cohen MR. Learning and attention reveal a general relationship between population activity and behavior. Science. 2018;359(6374):463–465.

8. Ratcliff R, McKoon G. The diffusion decision model: theory and data for two-choice decision tasks. Neural computation. 2008;20(4):873–922.

9. Gold JI, Shadlen MN. The neural basis of decision making. Annu Rev Neurosci. 2007;30(1):535–574.

10. Johansson RS, Flanagan JR. Coding and use of tactile signals from the fingertips in object manipulation tasks. Nature Reviews Neuroscience. 2009;10(5):345–359.

11. Weber AI, Saal HP, Lieber JD, Cheng JW, Manfredi LR, Dammann III JF, et al. Spatial and temporal codes mediate the tactile perception of natural textures. Proceedings of the National Academy of Sciences. 2013;110(42):17107–17112.

12. Boundy-Singer ZM, Saal HP, Bensmaia SJ. Speed invariance of tactile texture perception. Journal of neurophysiology. 2017;118(4):2371–2377.

13. Lezkan A, Drewing K. Processing of haptic texture information over sequential exploration movements. Attention, Perception, & Psychophysics. 2018;80(1):177–192.

14. Pei YC, Bensmaia SJ. The neural basis of tactile motion perception. Journal of Neurophysiology. 2014;112(12):3023–3032. doi:10.1152/jn.00391.2014.

15. Sherrick CE, Rogers R. Apparent haptic movement. Perception & Psychophysics. 1966;1(3):175–180.

16. Kirman JH. Tactile apparent movement: the effects of number of stimulators. Journal of Experimental Psychology. 1974;103(6):1175.

17. Kirman JH. Tactile apparent movement: The effects of interstimulus onset interval and stimulus duration. Perception & Psychophysics. 1974;15(1):1–6.

18. Geldard FA, Sherrick CE. The cutaneous “rabbit”: a perceptual illusion. Science. 1972;178(4057):178–179.

19. Geldard FA, Sherrick CE. The cutaneous saltatory area and its presumed neural basis. Perception & Psychophysics. 1983;33(4):299–304.

20. Gardner EP, Palmer CI. Simulation of motion on the skin. I. Receptive fields and temporal frequency coding by cutaneous mechanoreceptors of OPTACON pulses delivered to the hand. Journal of Neurophysiology. 1989;62(6):1410–1436.

21. Gardner EP, Palmer CI. Simulation of motion on the skin. II. Cutaneous mechanoreceptor coding of the width and texture of bar patterns displaced across the OPTACON. Journal of neurophysiology. 1989;62(6):1437–1460.

22. Olausson H, Norrsell U. Observations on human tactile directional sensibility. The Journal of physiology. 1993;464(1):545–559.

23. Olausson H, Wessberg J, Kakuda N. Tactile directional sensibility: peripheral neural mechanisms in man. Brain research. 2000;866(1-2):178–187.

24. Pei YC, Hsiao SS, Craig JC, Bensmaia SJ. Shape invariant coding of motion direction in somatosensory cortex. PLoS biology. 2010;8(2):e1000305.

25. Pei YC, Hsiao SS, Craig JC, Bensmaia SJ. Neural mechanisms of tactile motion integration in somatosensory cortex. Neuron. 2011;69(3):536–547.

26. Pack CC, Bensmaia SJ. Seeing and feeling motion: canonical computations in vision and touch. PLoS Biology. 2015;13(9):e1002271.

27. Mackevicius EL, Best MD, Saal HP, Bensmaia SJ. Millisecond precision spike timing shapes tactile perception. Journal of Neuroscience. 2012;32(44):15309–15317.

28. Kuroki S, Watanabe J, Nishida S. Neural timing signal for precise tactile timing judgments. Journal of Neurophysiology. 2016;115(3):1620–1629.

29. Kuroki S, Nishida S. Human tactile detection of within-and inter-finger spatiotemporal phase shifts of low-frequency vibrations. Scientific Reports. 2018;8(1):1–10.

30. Kuroki S, Nishida S. Motion direction discrimination with tactile random-dot kinematograms. i-perception. 2021;12(2):20416695211004620.

31. Birznieks I, McIntyre S, Nilsson HM, Nagi SS, Macefield VG, Mahns DA, et al. Tactile sensory channels over-ruled by frequency decoding system that utilizes spike pattern regardless of receptor type. Elife. 2019;8:e46510.

32. Prsa M, Morandell K, Cuenu G, Huber D. Feature-selective encoding of substrate vibrations in the forelimb somatosensory cortex. Nature. 2019;567(7748):384–388.

33. Prsa M, Kilicel D, Nourizonoz A, Lee KS, Huber D. A common computational principle for vibrotactile pitch perception in mouse and human. Nature communications. 2021;12(1):1–8.

34. Adibi M. Continuous vibration-driven virtual tactile motion perception across fingertips. Sensors. 2025;25(18):5918.

35. Rezaei E, Adibi M. Temporal Phase Differences Encode Tactile Motion from Static Vibrotactile Inputs. bioRxiv. 2026; p. 2026–01.

36. Kiani R, Shadlen MN. Representation of confidence associated with a decision by neurons in the parietal cortex. Science. 2009;324(5928):759.

37. Yeung N, Summerfield C. Metacognition in human decision-making: confidence and error monitoring. Philosophical Transactions of the Royal Society B: Biological Sciences. 2012;367(1594):1310.

38. Maniscalco B, Lau H. A signal detection theoretic approach for estimating metacognitive sensitivity from confidence ratings. Consciousness and cognition. 2012;21(1):422–430.

39. Meyniel F, Sigman M, Mainen ZF. Confidence as Bayesian probability: From neural origins to behavior. Neuron. 2015;88(1):78–92.

40. Balsdon T, Wyart V, Mamassian P. Confidence controls perceptual evidence accumulation. Nature communications. 2020;11(1):1753.

41. Young EM, Gueorguiev D, Kuchenbecker KJ, Pacchierotti C. Compensating for fingertip size to render tactile cues more accurately. IEEE transactions on haptics. 2020;13(1):144–151.

42. Treutwein B. Adaptive psychophysical procedures. Vision research. 1995;35(17):2503–2522.

43. Strauß J, Stritih-Peljhan N. Vibration detection in arthropods: Signal transfer, biomechanics and sensory adaptations. Arthropod Structure & Development. 2022;68:101167.

44. Mountcastle V, LaMotte R, Carli G. Detection thresholds for stimuli in humans and monkeys: Comparison with threshold events in mechanoreceptive afferent nerve fibers innervating the monkey hand. Journal of Neurophysiology. 1972;35(1):122–136.

45. Freeman AW, Johnson KO. Cutaneous mechanoreceptors in macaque monkey: temporal discharge patterns evoked by vibration, and a receptor model. The Journal of physiology. 1982;323(1):21–41.

46. Johansson RS, Landstro U, Lundstro R, et al. Responses of mechanoreceptive afferent units in the glabrous skin of the human hand to sinusoidal skin displacements. Brain research. 1982;244(1):17–25.

47. Bell J, Bolanowski S, Holmes MH. The structure and function of Pacinian corpuscles: a review. Progress in neurobiology. 1994;42(1):79–128.

48. Zimmerman A, Bai L, Ginty DD. The gentle touch receptors of mammalian skin. Science. 2014;346(6212):950–954.

49. Hollins EARM. A ratio code for vibrotactile pitch. Somatosensory & motor research. 1998;15(2):134–145.

50. Adibi M, Arabzadeh E. A comparison of neuronal and behavioral detection and discrimination performances in rat whisker system. Journal of Neurophysiology. 2011;105(1):356.

51. Adibi M, Diamond ME, Arabzadeh E. Behavioral study of whisker-mediated vibration sensation in rats. Proceedings of the National Academy of Sciences. 2012;109(3):971–976.

52. Adibi M. Whisker-mediated touch system in rodents: from neuron to behavior. Frontiers in Systems Neuroscience. 2019;13:40.

53. Braddick OJ. Low-level and high-level processes in apparent motion. Philosophical Transactions of the Royal Society of London B, Biological Sciences. 1980;290(1038):137–151.

54. Marr D. Vision: A computational investigation into the human representation and processing of visual information. MIT press; 2010.

55. Cavanagh P. Attention-based motion perception. Science. 1992;257(5076):1563–1565.

56. Lu ZL, Sperling G. Three-systems theory of human visual motion perception: review and update. Journal of the Optical Society of America A. 2001;18(9):2331–2370.

57. Hirsh IJ. Auditory perception of temporal order. The Journal of the Acoustical Society of America. 1959;31(6):759–767.

58. Bregman AS. Auditory scene analysis: The perceptual organization of sound. MIT press; 1994.

59. Alain C, Arnott SR, Hevenor S, Graham S, Grady CL. “What” and “where” in the human auditory system. Proceedings of the national academy of sciences. 2001;98(21):12301–12306.

60. Fink M, Ulbrich P, Churan J, Wittmann M. Stimulus-dependent processing of temporal order. Behavioural processes. 2006;71(2-3):344–352.

61. Brown AD, Jones HG, Kan A, Thakkar T, Stecker GC, Goupell MJ, et al. Evidence for a neural source of the precedence effect in sound localization. Journal of neurophysiology. 2015;114(5):2991–3001.

62. Stecker GC. Temporal weighting functions for interaural time and level differences. V. Modulated noise carriers. The Journal of the Acoustical Society of America. 2018;143(2):686–695.

63. Hamilton LS, Edwards E, Chang EF. A spatial map of onset and sustained responses to speech in the human superior temporal gyrus. Current Biology. 2018;28(12):1860–1871.

64. Ullman S. The interpretation of visual motion. Massachusetts Inst of Technology Pr; 1979.

65. Hirsh IJ, Sherrick Jr CE. Perceived order in different sense modalities. Journal of experimental psychology. 1961;62(5):423.

66. Parise CV, Ernst MO. Multisensory integration operates on correlated input from unimodal transient channels. ELife. 2025;12:RP90841.

67. Ratcliff R. A theory of memory retrieval. Psychological review. 1978;85(2):59.

68. Bogacz R, Brown E, Moehlis J, Holmes P, Cohen JD. The physics of optimal decision making: a formal analysis of models of performance in two-alternative forced-choice tasks. Psychological review. 2006;113(4):700.

